# The Human LRRK2-R1441G Mutation Drives Age-Dependent Oxidative Stress and Mitochondrial Dysfunction in Dopaminergic Neurons

**DOI:** 10.1101/2025.06.16.660029

**Authors:** Yuanxin Chen, Lianteng Zhi, Shiquan Cui, Huaixin Wang, Chenbo Zeng, Hui Zhang

## Abstract

Mitochondrial dysfunction and oxidative stress are central to Parkinson’s disease (PD) pathogenesis, particularly affecting substantia nigra pars compacta (SNc) dopamine (DA) neurons. Here, we investigate how the R1441G mutation in leucine-rich repeat kinase 2 (LRRK2), a key genetic contributor to familial and sporadic PD, impacts mitochondrial function in midbrain DA neurons. Using a BAC transgenic mouse model overexpressing human LRRK2-R1441G, we crossed these mice with TH-mito-roGFP mice, enabling mitochondria-targeted redox imaging in DA neurons. The two-photon imaging of acute brain slices from 3-, 6-, and 10-month-old mice revealed a progressive elevated oxidative stress in SNc DA neurons and their striatal projections, accompanied with reduced respiratory complex activity and decline in mitochondrial health. Spatial transcriptomics via GeoMx Digital Spatial Profiler identified molecular changes linked to dysregulated mitochondrial uncoupling protein function and calcium homeostasis. These findings demonstrate age-dependent mitochondrial dysfunction in LRRK2-mutant SNc DA neurons, highlighting calcium channels and uncoupling proteins as potential therapeutic targets to slow PD progression.

---

Parkinson’s disease (PD) is a progressive neurodegenerative disorder characterized by the selective loss of dopaminergic (DA) neurons in the substantia nigra pars compacta (SNc) and the accumulation of α-synuclein pathology (Lashuel et al., 2002; Dauer and Przedborski, 2003; Surmeier, 2018). While both genetic and environmental factors contribute to PD, aging remains the greatest risk factor (Fearnley and Lees, 1991). However, the mechanisms by which age-related cellular dysfunctions drive PD pathogenesis remain poorly understood. Among the most implicated factors, mitochondrial dysfunction and oxidative stress have emerged as critical contributors to dopaminergic neurodegeneration.

Mutations in leucine-rich repeat kinase 2 (LRRK2) are the most common genetic cause of both familial and sporadic PD (Zimprich et al., 2004; West et al., 2005; Kluss et al., 2019). Among these, the R1441G mutation disrupts the GTPase function of LRRK2, leading to aberrant kinase activity and pathological consequences, including abnormal mitochondrial function, oxidative stress, and calcium dysregulation (Healy et al., 2008; Cookson, 2010; Wang et al., 2012; Cherra et al., 2013). Although mitochondrial dysfunction has been implicated in PD, whether and how the LRRK2-R1441G mutation directly contributes to mitochondrial abnormalities, oxidative stress, and neurodegeneration in an age-dependent manner remains unresolved.

Mitochondria are essential for ATP production, calcium buffering, and reactive oxygen species (ROS) regulation (Sheu et al., 2006; Csordás and Hajnóczky, 2009). In DA neurons, which exhibit high metabolic demands due to autonomous pacemaking activity, mitochondrial health is particularly critical. Dysfunctional mitochondria lead to energy deficits, excessive ROS production, and oxidative damage, all of which are known contributors to neuronal vulnerability in PD (Chan et al., 2009). Notably, mitochondrial oxidative stress can be exacerbated by dysregulated calcium homeostasis, as excessive cytosolic and mitochondrial calcium levels further impair mitochondrial respiration and trigger cell death pathways.

Here, we investigate the role of mitochondrial dysfunction and oxidative stress in LRRK2-R1441G-associated PD using a BAC transgenic mouse model that overexpresses human LRRK2-R1441G. This model exhibits progressive motor, neurochemical, and pathological deficits, closely mirroring human PD (Li et al., 2009). By crossing these mice with TH-mito-roGFP reporters and applying two-photon ratiometric imaging, we uncovered age-dependent mitochondrial impairment marked by early oxidative stress in striatal DA terminals and calcium overload driven by increased MCU expression. These deficits were mitigated by L-type calcium channel blockade, implicating calcium dysregulation as a driver of oxidative stress. Furthermore, we observed impaired mitochondrial membrane potential (MMP) flickering and reduced UCP4/UCP5 expression, exacerbating ROS accumulation (Kim-Han and Dugan, 2005; Guzman et al., 2010) was significantly impaired in hR1441G DA neurons. Spatial transcriptomics of the substantia nigra further confirmed the upregulation of redox stress pathways in LRRK2-R1441G DA neurons. Taking together, our study provides the first direct mechanistic link between hLRRK2-R1441G, mitochondrial oxidative stress, and calcium dysregulation in PD., highlighting UCPs, MCU, and L-type calcium channels as potential therapeutic targets in PD.

## Result

### LRRK2-hR1441G Mutation Disrupts Mitochondrial Morphology and Impairs Respiratory Function in Dopaminergic Terminals within the Striatum

Mutations in LRRK2 represent the most common genetic cause of both familial and sporadic Parkinson’s disease (PD). Li et al. (2009) reported age-dependent motor deficits in hLRRK2-R1441G mice, implicating this mutation in PD pathogenesis. Mitochondrial dysfunction is a key driver of dopaminergic neuron degeneration in the substantia nigra pars compacta (SNc), with striatal terminal loss preceding neuronal loss in the SNc. To determine whether mitochondrial dysfunction occurs specifically at dopaminergic terminals, we employed tyrosine hydroxylase (TH) immunostaining combined with electron microscopy (EM) to assess mitochondrial morphology. In 10-month-old hR1441G TG mice, mitochondria in TH-positive regions exhibited abnormal structural features, including cristae loss and irregular oval-shaped morphology (Fig. 1a, b). In contrast, wild-type (WT) mice displayed predominantly healthy mitochondria with regular tubular morphology in TH-positive terminals (Fig. 1c). Quantitative analysis of mitochondrial aspect ratio (i.e., major axis length divided by minor axis length) further revealed that mitochondria in hR1441G mice were closer to unity, suggesting increased mitochondrial fission (Fig. 1d). Despite these qualitative changes, the overall striatal distribution of healthy mitochondria did not differ significantly between hR1441G TG and WT mice. These findings indicate that the R1441G mutation leads to abnormal mitochondrial morphology only in dopaminergic terminals, which may contribute to early synaptic dysfunction in PD.

**Fig. 1.**
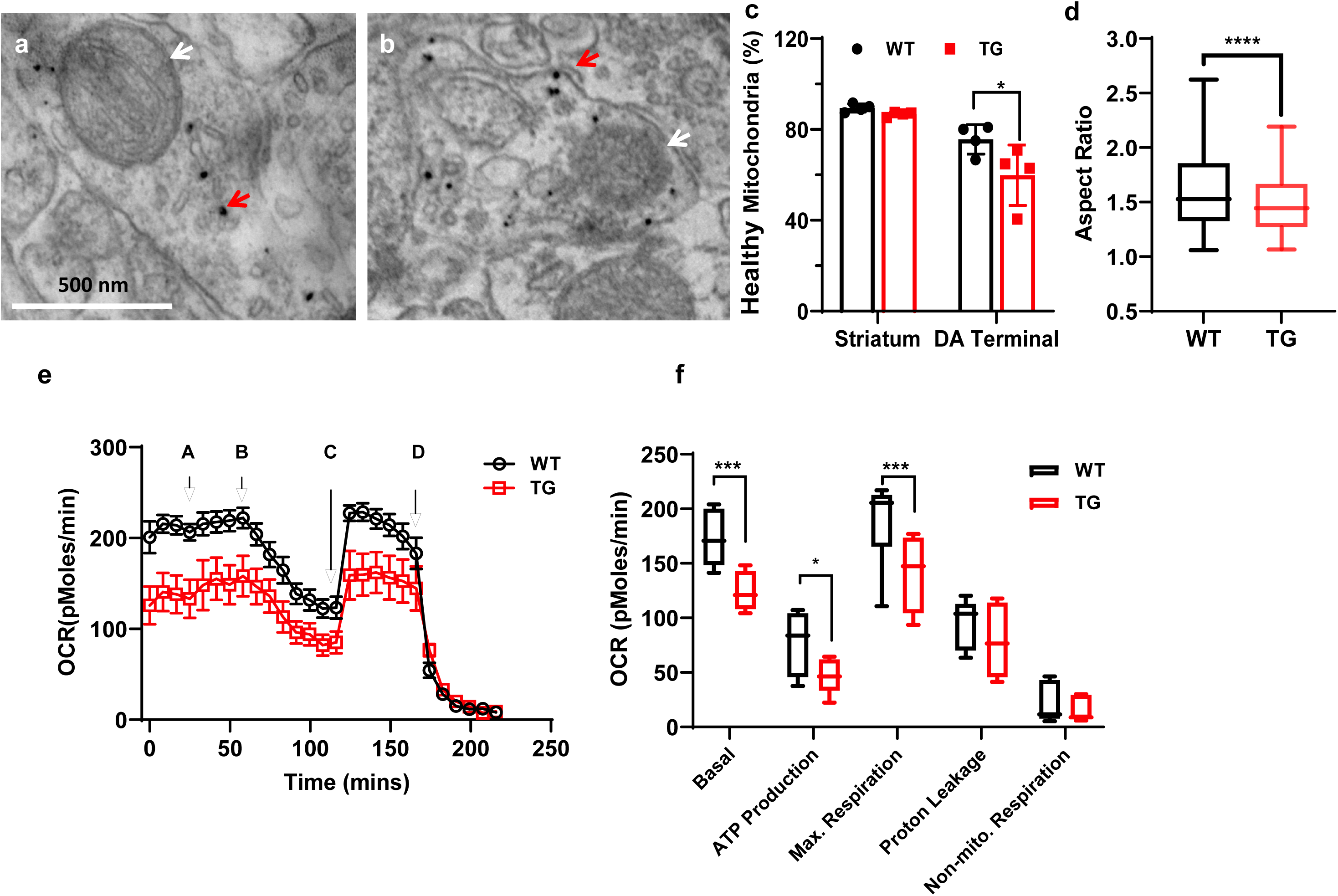
Mitochondria respiration in LRRK2 WT and R1441G TG striatum slices. **a, b,** Representative EM images of mitochondria after TH immunogold staining in striatal slices from age matched LRRK2 hR1441G WT and TG mice, respectively. Red arrows indicated tyrosine hydroxylase (TH)-positive signals within dopamine terminals. White arrows pointed to mitochondria located within or around the dopamine terminals. **c,** Quantification of mitochondrial density in the striatum and in proximity to dopaminergic (DA) terminals. Compared to WT controls, hR1441G TG mice displayed a significant increase in the proportion of unhealthy mitochondria near DA terminals (striatum: WT, *N* = 4, *n* = 1300; TG, *N* = 4, *n* = 1231; DA terminal: WT, *n* = 52; TG, *n* = 67, \**P* = 0.0225). **d,** Mitochondrial aspect ratio, an indicator of morphology, was significantly higher in WT mice compared to TG mice (unpaired Mann-Whitney test, \*\*\*\**P* < 0.0001). **e,** Acute striatal slices from WT and TG mice were sequentially treated with 10 mM pyruvate (A), 10 µM oligomycin (B), 10 µM FCCP (C), and 20 µM antimycin A (D). Real-time oxygen consumption rates (OCR), indicative of oxidative phosphorylation, were measured. **f,** LRRK2 TG mice exhibited a significant reduction in basal respiration, ATP production, and maximal respiration, while proton leakage and non-mitochondrial respiration remained unaltered (two-way ANOVA test, Basal, WT *vs* TG, \*\*\**P* = 0.0006; ATP production, WT *vs* TG \**P* = 0.0478; Max. Respiration, WT *vs* TG, \*\*\**P* = 0.0006; Proton Leakage, *P* = 0.5808; non-mitochondrial Respiration, *P* = 0.9972).

To assess whether the LRRK2-hR1441G mutation impairs mitochondrial respiration, we performed Seahorse respirometry on acutely prepared striatal slices sequentially treated with 10 mM pyruvate, 10 µM oligomycin, 10 µM FCCP, and 20 µM antimycin (*Zhi et al., 2018).* In 10-month-old hLRRK2-R1441G transgenic mice, we observed a significant reduction in basal oxygen consumption rate (OCR), ATP production, and maximal respiratory capacity (Fig. 1e, f; Extended Data Fig. 1). However, no significant differences in mitochondrial respiration were detected in younger (3- and 6-month-old) animals, suggesting a progressive decline in mitochondrial function with aging.

### hR1441G Mutation Induces Age-Dependent Mitochondrial Oxidative Stress in SNc Dopaminergic Neurons and Striatal Terminals

The autonomous pace-making activity of substantia nigra pars compacta (SNc) dopaminergic neurons regulate dopamine release, which is essential for striatal function. This process depends on ATP production via mitochondrial oxidative phosphorylation, which concurrently generates superoxide and reactive oxygen species (ROS), leading to mitochondrial oxidative stress.

To assess basal oxidative stress levels in the SNc and striatum of an LRRK2-hR1441G PD model, we crossed this model with TH-mito-roGFP mice, which express a mitochondrially targeted, redox-sensitive green fluorescent protein under the TH promoter, creating hLRRK2-R1441G/TH-mito-roGFP mice. Using two-photon laser scanning microscopy (2PLSM) combined with a ratiometric imaging approach (Fig. 2a-c; Extended Data Fig. 1–4; details in Supplementary Materials), we assessed mitochondrial redox states in dopaminergic neurons in acute brain slices. LRRK2-hR1441G mutant mice exhibited significantly increased mitochondrial oxidation levels in the SNc and striatum, particularly at 6 and 12 months of age (Fig. 2d–f; Extended Data Fig. 5–6). Notably, in 3-month-old LRRK2-hR1441G mice, oxidative stress was already elevated in striatal terminals, suggesting early mitochondrial dysfunction at presynaptic sites. This oxidative stress increase followed an age-dependent pattern. In SNc dopaminergic neurons, oxidation levels progressively increased from 3 to 6 months and further at 12 months. In contrast, in striatal terminals, oxidative stress rose significantly between 3 and 6 months but plateaued thereafter, with no further increase between 6 and 12 months in LRRK2-hR1441G mice. No significant differences in mitochondrial redox states were detected in the ventral tegmental area (VTA) between 6-month-old WT and LRRK2-hR1441G mice (Fig. 2g; Extended Data Fig. 7) (Mann-Whitney test, P = 0.8981).

**Fig. 2.**
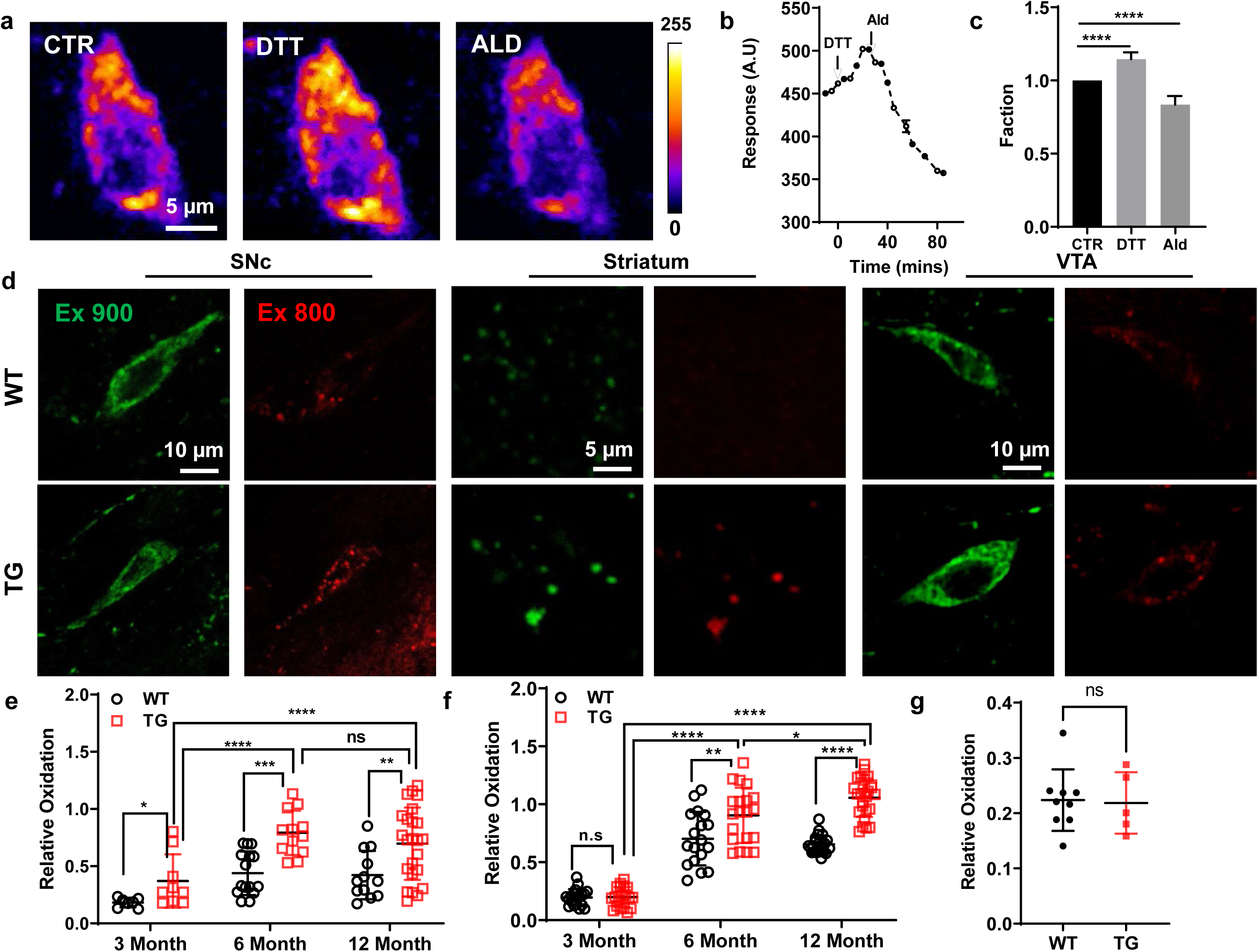
Aged-dependent oxidative stress of LRRK2 R1441G WT and TG animal model. **a,** Visualization of substantia nigra pars compacta (SNc) dopaminergic (DA) neurons expressing roGFP under different conditions: control, dithiothreitol (DTT), and aldehyde (Ald) treatments. **b,** Quantitative analysis of roGFP fluorescence intensity over time following sequential treatment with DTT and Ald. **c,** Statistical analysis of the effects of DTT and Ald on roGFP fluorescence intensity (*N* = 4, *n* = 6, Wilcoxon matched pairs signed rank test; CTR *vs* DTT, \**P* = 0.0313; CTR *vs* Ald, \**P* = 0.0313). **d,** Representative images of SNc DA neurons expressing roGFP in 6-month-old LRRK2 WT and TG mice under two excitation wavelengths (900 nm and 800 nm) and across different brain regions (striatum, SNc, and VTA). **e,** Oxidative stress levels in the striatum also showed an age-dependent increase, with hLRRK2 R1441G TG mice demonstrating elevated oxidative stress at 3 month (\**P* = 0.0140), 6 months (\*\*\**P* = 0.0008) and over 12 months (\*\**P* = 0.0051) compared to WT controls (3 pairs for 3-month-old WT, *n* = 10; TG, *n* = 7; 4 pairs for 6-month-old WT, *n* = 11; TG, *n* = 8; 3 pairs for over 12-month-old WT, *n* = 12; TG, *n* = 20). **f,** Oxidative stress levels in SNc DA neurons increased with age, with hLRRK2 TG mice exhibiting significantly higher oxidative stress at 6 months (\*\**P* = 0.005) and 12 months (\*\*\*\**P* < 0.0001) compared to WT controls (3 pairs; DA neurons: 3-month-old WT, *n* = 18; TG, *n* = 21; 6-month-old WT, *n* = 18; TG, *n* = 21; 12-month-old WT, *n* = 24; TG, *n* = 25). **g,** No significant difference in oxidative stress was observed in VTA DA neurons between 6-month-old LRRK2 WT and TG mice (3 pairs; DA neurons: WT, *n* = 9; TG, *n* = 5; Mann-Whitney test, *P* = 0.8981).

These findings demonstrate that the LRRK2-hR1441G mutation induces region-specific and age-dependent increases in mitochondrial oxidative stress, particularly in the SNc and dorsal striatum, but not in the VTA. This suggests that SNc DA neurons and their striatal projections are more vulnerable to oxidative stress, which may contribute to their selective degeneration in PD.

### Cytosolic and mitochondrial Ca^2+^ are overloaded in hLRRK2 R1441G mice

To investigate whether the elevated oxidative stress in LRRK2-hR1441G TG mice results from altered calcium homeostasis, we injected AAV-GCaMP6 under the control of the tyrosine hydroxylase (TH) promoter into the substantia nigra pars compacta (SNc) to selectively target dopaminergic neurons. Quantification of cytosolic calcium levels revealed a significant increase in LRRK2-hR1441G TG mice, with fluorescence intensity normalized to its maximum value rising from 19.71% in WT mice to 75.75% in TG mice, indicating a substantial elevation in cytosolic calcium induced by the LRRK2-hR1441G mutation (Fig. 3a-b; Mann-Whitney test, P = 0.0012; WT: N = 3, n = 7; TG: N = 3, n = 6).

**Fig. 3.**
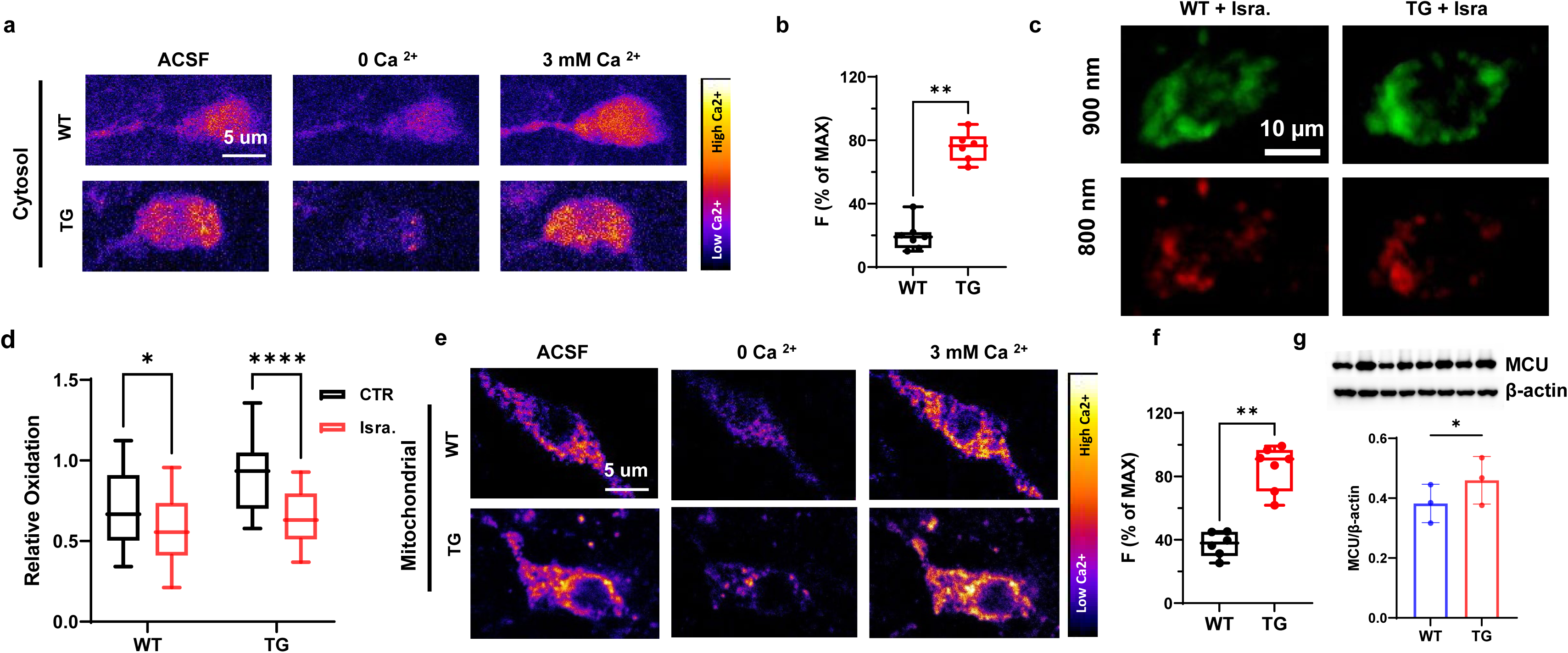
LRRK2 R1441G Mutation Elevates Cytosolic and Mitochondrial Calcium Levels in 5-Month-Old Mice a, Pseudo-colored representative images of cytosolic calcium levels in striatal slices from LRRK2 WT and hR1441G mice under varying perfusion conditions: regular artificial cerebrospinal fluid (ACSF), ACSF with 0 mM Ca² + EGTA + 1 μM ionomycin, and ACSF with 3 mM Ca² + 1 μM ionomycin. **b,** Quantification of cytosolic calcium levels revealed that the LRRK2 hR1441G mutation significantly increased cytosolic calcium concentration compared to WT mice (Mann-Whitney test, \*\**P* = 0.0012; WT: *N* = 3, *n* = 7; TG: *N* = 3, *n* = 6). c, Representative images illustrating the effect of the L-type calcium channel blocker Isradipine on redox levels in SNc dopaminergic neurons. **d**, Isradipine treatment significantly reduced redox levels in SNc dopaminergic neurons (two-way ANOA, WT *vs* WT + Isra., \**P* = 0.0427, TG *vs* TG + TG + Isra., \*\*\*\**P* < 0.0001). **e,** Pseudo-colored representative images of mitochondrial calcium levels in LRRK2 WT and TG mice under different perfusion conditions: regular ACSF, ACSF with 0 mM Ca² + EGTA + 1 μM ionomycin, and ACSF with 3 mM Ca² + 1 μM ionomycin. **f**, Quantification of mitochondrial calcium indicated that LRRK2 R1441G mutation significantly increased mitochondrial calcium concentration (Mann Whitney test, \*\**P* = 0.0012; WT, *N* = 3 *n* = 6; TG, *N* = 4, *n* = 7). **g**, Top: Representative Western blot analysis of mitochondrial calcium uniporter (MCU) expression in the substantia nigra (SN) of LRRK2 WT and TG mice. Bottom: Quantitative analysis revealed a significant elevation in MCU expression levels in LRRK2 hR1441G mice (3 pairs, Mann-Whitney test, \**P* = 0.0138).

To determine whether ROS elevation was linked to L-type calcium channel activity, we administered isradipine, an L-type calcium channel inhibitor, to LRRK2-hR1441G mice. Redox level measurements in SNc dopaminergic neurons demonstrated that isradipine significantly reduced ROS levels, supporting a direct link between calcium dysregulation and oxidative stress (Fig. 3c-d; two-way ANOVA; WT, P = 0.0047; TG, P < 0.0001).

To further assess mitochondrial calcium levels, we injected AAV vectors expressing a mitochondria-targeted calcium indicator under the TH-mito-matrix promoter into the SNc of WT and TG mice. The hR1441G mutation significantly increased mitochondrial calcium concentrations (Fig. 3e-f; Mann-Whitney test, P = 0.0012; WT: N = 3, n = 6; TG: N = 4, n = 7). Western blot analysis further revealed upregulation of the mitochondrial calcium uniporter (MCU), a key regulator of calcium influx into the mitochondria, in hR1441G TG mice, supporting a role for MCU-mediated calcium overload in mitochondrial dysfunction (Fig. 3g).

Collectively, these findings demonstrate that calcium overload, driven by increased mitochondrial calcium uptake and elevated MCU expression, contributes to excessive ROS production in LRRK2-hR1441G mutant dopaminergic neurons, establishing a mechanistic link between calcium dysregulation and oxidative stress in PD pathogenesis.

### Mitochondrial membrane potential (MMP) transient is linked to oxidative stress

The mitochondrial membrane potential (MMP) plays a critical role in regulating ATP production and reactive oxygen species (ROS) generation (Guzman et al., 2010). To investigate whether elevated oxidative stress in hR1441G mice correlates with alterations in MMP, we used tetramethylrhodamine methyl ester (TMRM, 2–4 µM), a dye that selectively accumulates within mitochondria to measure MMP. Since TMRM labeling is not cell-type specific, we generated hLRRK2-R1441G mice expressing TH-GFP to selectively identify dopaminergic neurons among TMRM-labeled populations (Extended Data Fig. 8). We observed periodic TMRM signals of DA neurons in both WT and TG mice, indicating transient changes in MMP (Fig. 4a; Extended Data Video 1). However, the percentage, amplitude and the frequency of MMP transients in SNc dopaminergic neurons in TG mice was significantly reduced, compared to WT mice (Fig. 4b-c). Notably, MMP flickering, indicative of MMP depolarization events, was observed more frequently in dopaminergic neurons of WT mice (Fig. 4c). The frequency of MMP transients exhibited an age-dependent decline, decreasing progressively from 3 to 6 to 12 months in both genotypes. However, TG mice showed significantly reduced MMP transient frequencies at 6 and 12 months compared to WT mice, with no significant differences observed at 3 months (Extended Data Fig. 9).

**Fig. 4.**
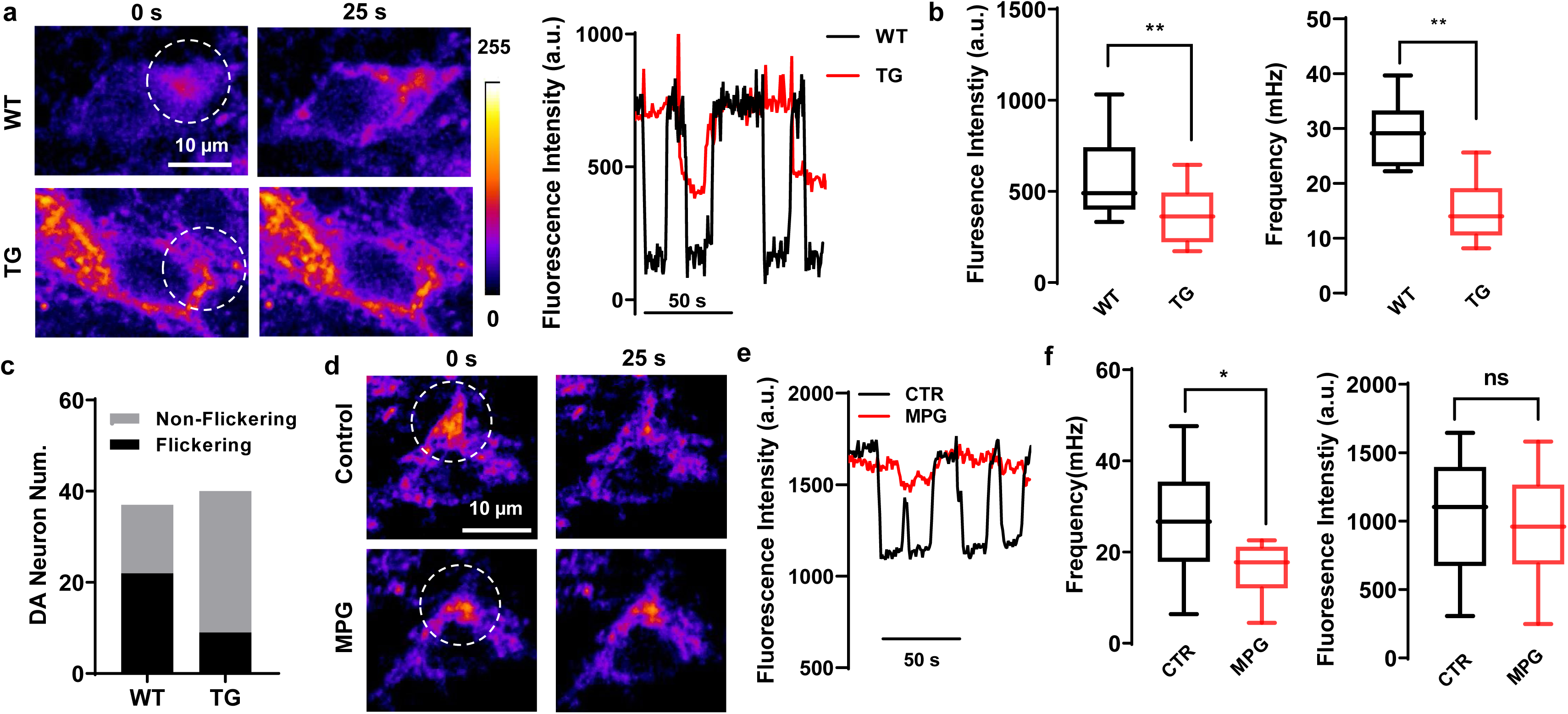
Mitochondrial membrane potential (MMP) transient is linked to oxidative stress. **a**, Left, Representative images of mitochondria in SNc dopaminergic neurons in brain slices labeled with TMRM at the different time (0 sec and 25 sec) from LRRK2 WT and hR1441G mice. Right: Time series of TMRM fluorescence intensity in a region of interest (ROI, white circle) from the left panel, illustrating the difference of MMP dynamics between LRRK2 WT and TG. **b**, **Left**, Box plot showing a significant reduction in MMP amplitude in LRRK2 TG mice compared to WT (Mann-Whitney test, \*\**P* = 0.0073). **Right**, Box plot demonstrated that flickering frequency was significantly higher in LRRK2 TG mice than in WT (Mann-Whitney test, \*\**P* = 0.0011). **c**, Bar graph summarizing the number of SNc dopaminergic neurons with observable MMP flickering events, indicating a lower number of neurons exhibiting flickering in LRRK2 TG mice. **d**, Representative images of SNc dopaminergic neurons labeled with TMRM before and after the bath treatment of an antioxidant, N-(2-mercaptopropionyl)-glycine (MPG). **e**, Time series of TMRM fluorescence intensity before and after MPG application, showing a decrease in flickering events following MPG treatment. **f**, Box plot summarizing flickering frequency (left) and amplitude (right) of flickering events after MPG treatment (*N* = 3, *n* = 9), indicating that MPG effectively mitigates flickering events associated with oxidative stress (Mann-Whitney test, \**P* = 0.0260).

To determine whether oxidative stress underlies MMP flickering, we treated brain slices with N-(2-mercaptopropionyl) glycine (MPG), a cell-permeable antioxidant. MPG treatment reduced MMP flickering and decreased MMP amplitudes (Fig. 4d-f; Extended Data Video 2).

These findings collectively suggest that mitochondrial flickering is associated with elevated oxidative stress and that the R1441G mutation disrupts MMP homeostasis in an age- and oxidative stress-dependent manner.

### Uncoupling protein is linked to mitochondrial flickering events

Calcium overload has been shown to elevate redox levels. To determine whether calcium activity influences MMP dynamics, we applied isradipine, an L-type calcium channel antagonist, to assess the relationship between calcium influx and MMP flickering events in SNc dopaminergic neurons. Isradipine treatment significantly diminished MMP transients and reduced MMP amplitudes (Fig. 5a-d; Extended Data Video 3), indicating that MMP flickering events are tightly associated with calcium influx.

**Fig. 5.**
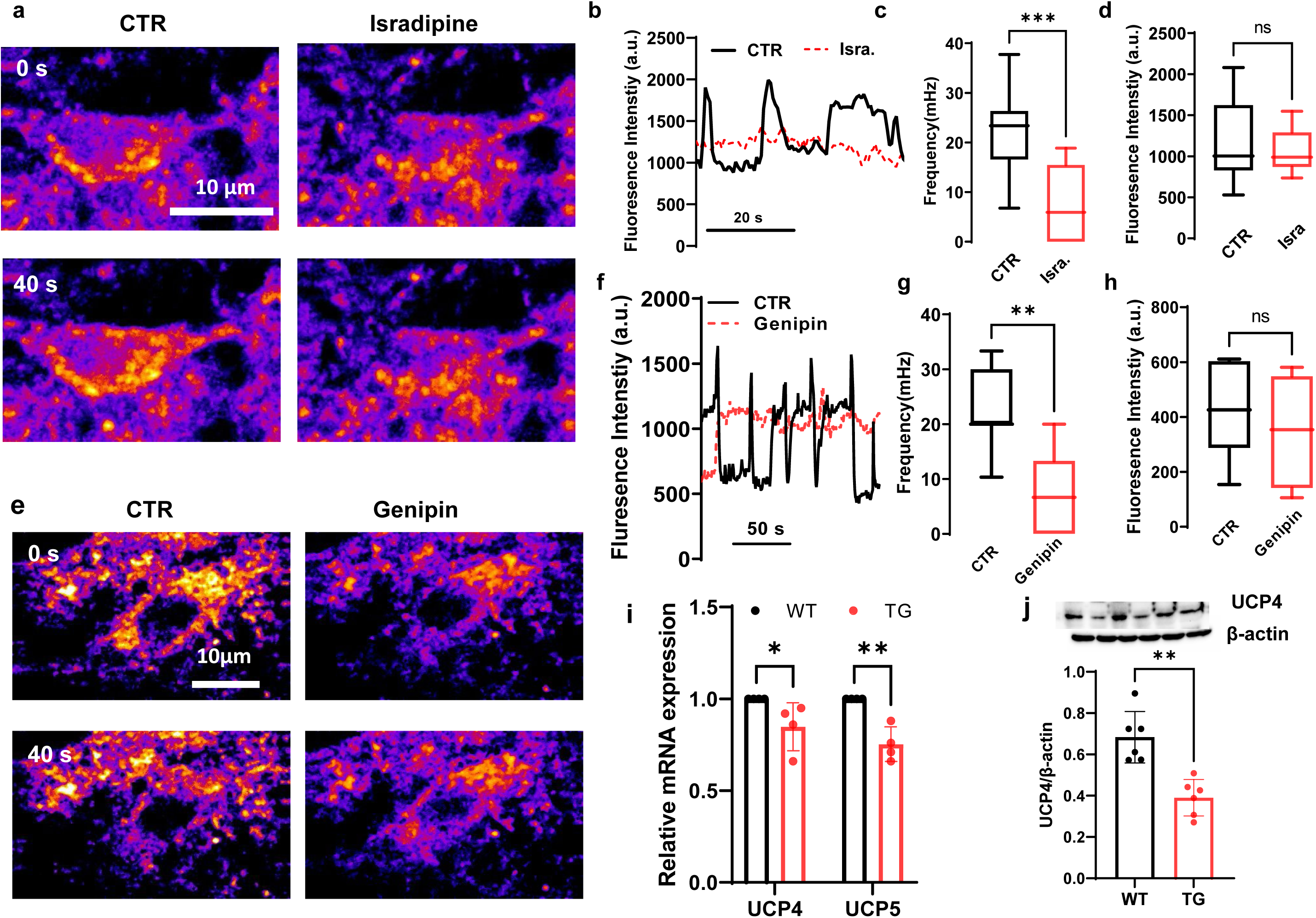
Uncoupling protein is linked to mitochondrial flickering events. **a**, Representative TMRM fluorescence images of SNc dopaminergic neurons before and after treatment with Isradipine. **b**, Isradipine significantly reduces mitochondrial flickering frequency. c and d, Quantitative analysis of Isradipine’s effects on flickering frequency and amplitude in SNc dopaminergic neurons, demonstrating a significant reduction in flickering events following treatment (*N* = 4, *n* = 8; Mann-Whitney test, \*\*\**P* = 0.0010). **e**, **f**, Representative TMRM fluorescence time-series images of SNc dopaminergic neurons before and after genipin treatment, respectively, indicating a decrease in mitochondrial flickering frequency after genipin application. **g**, **h**, Quantitative analysis of genipin’s effects on flickering frequency and amplitude in SNc dopaminergic neurons, respectively, showing a significant reduction in flickering events following treatment (*N* = 6, *n* = 10; Mann-Whitney test, \*\**P* = 0.0052). **i**, mRNA expression analysis of uncoupling proteins (UCPs) reveals that UCP4 and UCP5 levels are significantly reduced in TG mice compared to WT (4 pairs, two-way ANOVA, WT vs TG, UCP4, \**P* = 0.0397, UCP5, \*\**P* = 0.0019). **j**, Top: Representative Western blot showing UCP4 expression levels in the substantia nigra (SN) of LRRK2 hR1441G mice. Bottom: Quantitative analysis indicated a significant reduction in UCP4 expression in the SN of LRRK2 hR1441G mice compared to WT (4 pairs; Mann-Whitney test, \*\**P* = 0.0022).

Uncoupling proteins (UCPs) are ion channels in the inner mitochondrial membrane that mitigate oxidative stress by dissipating the proton gradient across the membrane. In the brain, UCPs reduce superoxide production, serving as a negative-feedback mechanism to limit reactive oxygen species (ROS) generation (Kim-Han and Dugan, 2005; Guzman et al., 2010). Elevated oxidative stress can increase UCP activation, thereby reducing the IMM potential. To investigate whether MMP flickering is influenced by UCP activity, we applied genipin, a UCP inhibitor, to brain slices containing SNc neurons exhibiting MMP flickering. Genipin significantly attenuated the frequency of MMP flickering events (Fig. 5e-h; Extended Data Video 4), suggesting that UCP activity directly contributes to MMP dynamics within the mitochondrial IMM.

To examine whether the LRRK2 hR1441G mutation affects UCP expression, we isolated mRNA from mitochondria from SNc dopaminergic neurons using laser microdissection and quantified mRNA levels of UCP components. Compared to WT mice, hR1441G TG mice exhibited significantly reduced mRNA levels of UCP4 and UCP5 (Fig. 5i). This finding was further corroborated by western blot analysis, which revealed a decrease in UCP4 protein expression in hR1441G TG mice (Fig. 5j).

These results suggest that the hR1441G mutation impairs UCP expression, contributing to altered mitochondrial membrane potential and oxidative stress.

### Digital spatial profiling (DSP) reveals hLRRK2 R1441G mutation induced elevation of ROS in dopaminergic neuron in SNc

To elucidate the molecular pathways disrupted by the *LRRK2* hR1441G mutation, we performed GeoMx digital spatial profiler (DSP) analysis on dopaminergic (DA) neurons in the SNc, comparing gene expression profiles between hR1441G transgenic (TG) and wild-type (WT) mice. Uniform Manifold Approximation and Projection (UMAP) analysis confirmed clear segregation of DA neurons from other types of cells: microglia and astrocytes in the SNc (Fig. 6a-b). The analysis identified 19 significantly upregulated and 9 downregulated genes in mutant DA neurons (Fig. 6c). Volcano plots further highlighted the differentially expressed genes (DEGs) in the transcriptional profiles caused by the LRRK2 hR1441G mutation (Fig. 6d).

**Fig. 6.**
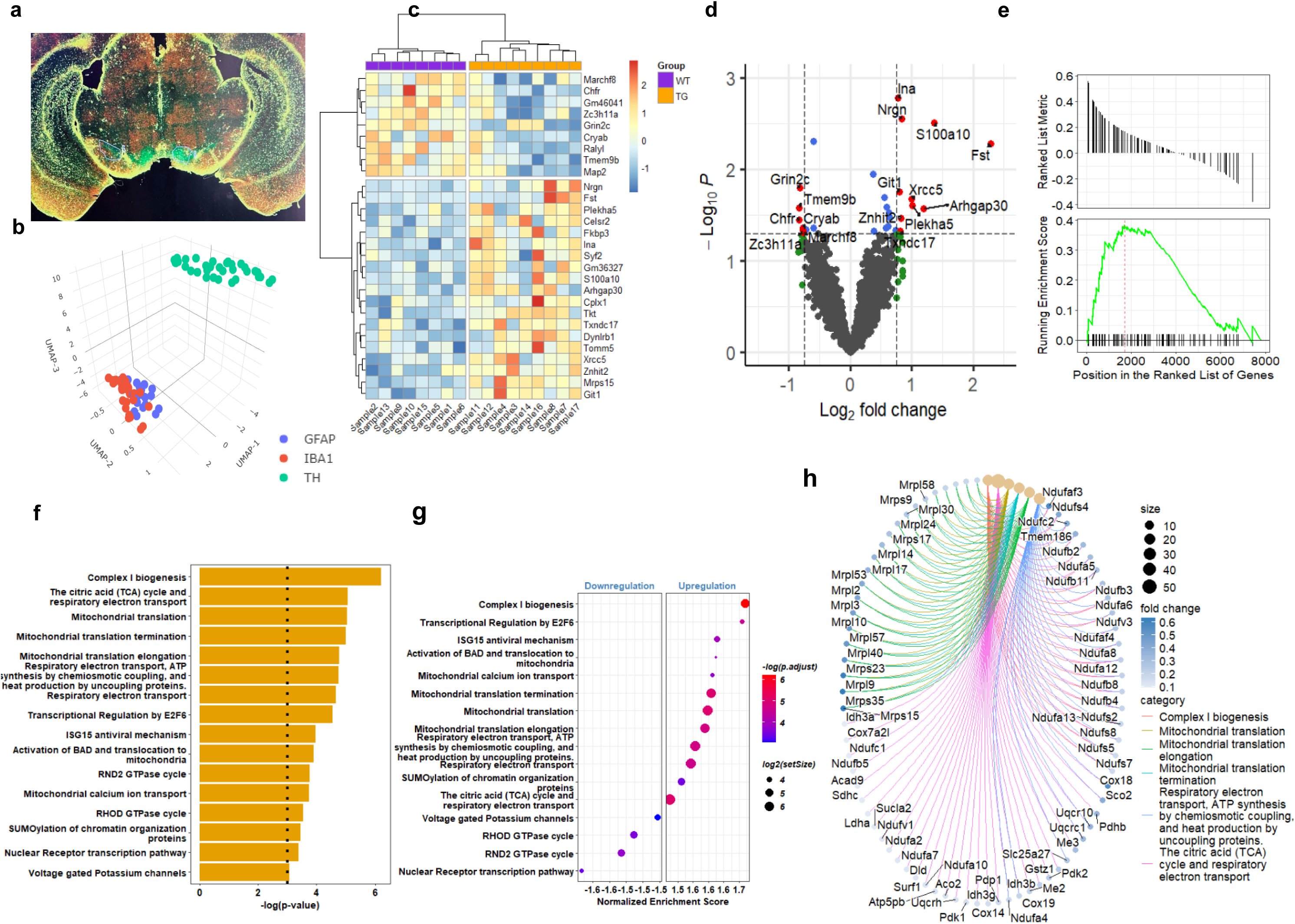
Digital Spatial Profiling of Dopaminergic (DA) Neurons in the Substantia Nigra (SN) **a**, Representative coronal brain sections displaying the spatial distribution of three cell markers: tyrosine hydroxylase (TH), GFAP, and IBA1, used for digital spatial profiling in the brain region SNc. **b**, UMAP clustering reveals distinct cell types. **c,** Heatmap displaying significant differentially expressed genes (DEGs) in DA neurons between LRRK2 TG and WT mice within the SN region. **d**, Volcano plots visualizing DEGs distribution in DA neurons between LRRK2 TG and WT mice, with Log_2_ fold changes plotted against -Log_10_ of the adjusted p-value for each gene. **e**, Enrichment plot showing upregulation of gene sets in DA neurons from LRRK2 TG mice relative to WT, associated with terms: respiratory electron transport, ATP synthesis via chemiosmotic coupling, and heat production by uncoupling proteins. **f**, Bar plot depicting the top 16 enrichment terms derived from the DEGs of DA neurons, ranked by statistical significance. **g**, Gene set enrichment analysis (GSEA) in DA neurons is presented as a hierarchically clustered heatmap. The color key represents the -Log10 p-value for the enrichment score, while the circle size reflects the number of leading genes associated with each enrichment term. **h**, CNETplot visualizing the complex associations between genes and significantly enriched GO terms in DA neurons. Node size corresponds to the statistical significance (adjusted p-value) of the term, while gene node colors indicate their fold change values. Edges denote the relationship between genes and their associated enrichment terms.

**Fig. 7.**
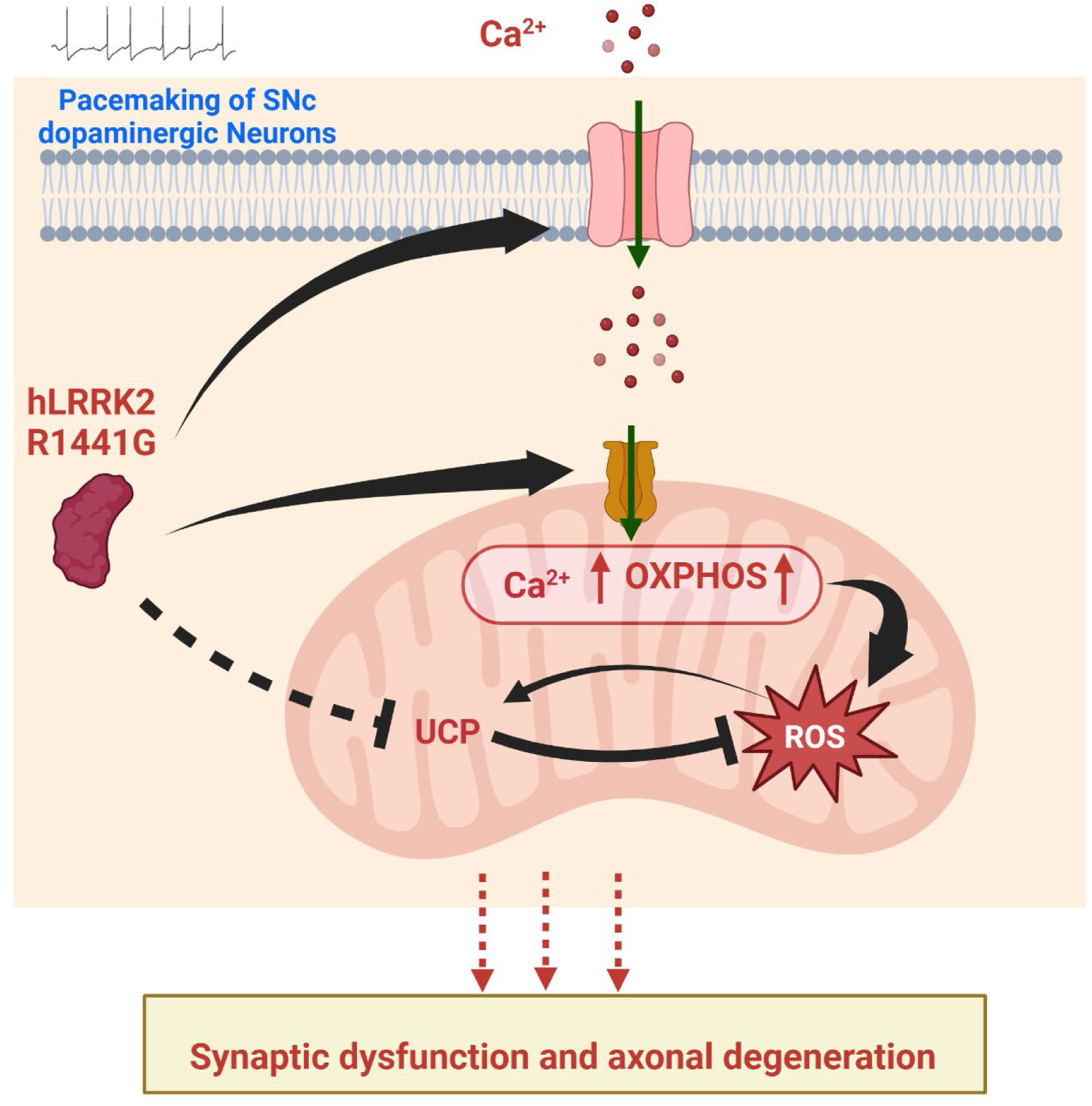
Summary of findings linking LRRK2 mutation to elevated mitochondrial oxidative stress. The LRRK2 R1441G mutation contributes to heightened mitochondrial oxidative stress in dopaminergic neurons within the substantia nigra pars compacta (SNc). This mutation facilitates increased cytosolic calcium influx and enhances mitochondrial calcium accumulation, mediated by the upregulation of mitochondrial calcium uniporter (MCU) protein. Elevated mitochondrial calcium concentrations drive an upregulation of oxidative phosphorylation (OXPHOS), leading to increased production of reactive oxygen species (ROS). Furthermore, the LRRK2 R1441G mutation downregulates uncoupling proteins (UCPs), impairing the degradation and clearance of ROS. This imbalance exacerbates oxidative stress, promoting synaptic dysfunction and axonal degeneration in SNc dopaminergic neurons. These pathological processes highlight the critical role of LRRK2 R1441G in mitochondrial dysfunction and neurodegeneration.

Functional enrichment analysis of DEGs revealed significant upregulation of pathways related to complex I biogenesis, mitochondrial calcium ion transport, respiratory electron transport, and heat production mediated by uncoupling proteins (UCPs) in LRRK2 hR1441G mutant DA neurons (Fig. 6e). Among the top 16 significantly enriched terms, the majority were associated with elevated redox activity, including “Complex I biogenesis,” “The citric acid (TCA) cycle and respiratory electron transport,” and “Oxidative phosphorylation – Mus musculus (house mouse)” (Fig. 6f-g; Extended Data Fig. 10a-c). Gene association analyses further revealed that most DEGs contributing to elevated ROS generation were related to mitochondrial function in origin (Fig. 6h; Extended Data Fig. 10d-e). These findings underscore that the LRRK2 hR1441G mutation profoundly disrupts mitochondrial function and exacerbates pathways linked to ROS production in SNc DA neurons.

Our findings provide the first direct evidence linking oxidative stress to the LRRK2-hR1441G mutation. Specifically, this mutation disrupts calcium homeostasis, leading to elevated cytosolic and mitochondrial calcium levels, increased basal oxidative stress, and a reduction in mild uncoupling events, likely due to UCP dysfunction or decreased expression. These results suggest potential new therapeutic strategies for LRRK2-associated PD, including enhancing UCP expression or targeting UCP activity with agonists, inhibiting MCU functions, mitigating oxidative stress and restore mitochondrial function, in addition to blocking L-type channels. Mitochondrial dysfunction and oxidative stress are recognized as key drivers of dopaminergic neurodegeneration in PD. However, the mechanisms underlying LRRK2-mediated mitochondrial impairment remain incompletely understood. Here, we provide direct evidence that the LRRK2-hR1441G mutation induces mitochondrial oxidative stress, disrupts calcium homeostasis, and impairs mitochondrial function in an age-dependent manner. Our findings highlight a mechanistic link between pathogenic LRRK2 mutation, oxidative stress, and calcium dysregulation, offering potential targets for therapeutic intervention in PD.

## Discussion

Our study demonstrates that LRRK2-hR1441G mice exhibit progressive mitochondrial oxidative stress in SNc DA neurons and their striatal projections. Using two-photon ratiometric imaging of TH-mito-roGFP mice, we show that oxidative stress is elevated at dopaminergic terminals as early as 3 months of age, preceding significant neurite loss. This finding supports the hypothesis that presynaptic mitochondrial dysfunction is an early event in PD and may contribute to progressive neurodegeneration.

We also observed that mitochondrial oxidative stress increased progressively in SNc DA neurons from 3 to 12 months of age, whereas oxidative stress in striatal terminals plateaued after 6 months. This suggests that early presynaptic mitochondrial dysfunction may initiate the disease process, with somatic mitochondrial stress accumulating later, consistent with axon-first degeneration models of PD (Burke and O’Malley, 2013; Hattori and Mizuno, 2015). In contrast, no significant oxidative stress changes were observed in the VTA, reinforcing the selective vulnerability of SNc DA neurons to mitochondrial dysfunction.

A key finding of our study is that LRRK2-hR1441G mutant DA neurons exhibit calcium overload, which contributes to mitochondrial dysfunction. Cytosolic calcium levels were significantly elevated, and mitochondrial calcium uptake was markedly increased, correlating with increased expression of the MCU. Pharmacological inhibition of L-type calcium channels with isradipine significantly reduced oxidative levels, implicating excessive calcium influx as a key driver of oxidative stress.

These findings support a pathogenic model in which LRRK2 mutations impair calcium homeostasis, leading to mitochondrial calcium overload, respiratory dysfunction, and ROS accumulation. This is particularly relevant given evidence that Ca² overload sensitizes DA neurons to oxidative damage, driving mitochondrial permeability transition pore (mPTP) opening and cell death. Our results further implicate UCP dysfunction in exacerbating oxidative stress, as LRRK2-hR1441G DA neurons exhibited reduced UCP4/UCP5 expression, impairing mitochondrial membrane potential flickering, a mechanism that protects against excessive ROS buildup.

Digital spatial profiling (DSP) added an important molecular dimension to our findings, revealing that dopaminergic neurons in TG mice exhibit widespread upregulation of genes involved in oxidative phosphorylation (OXPHOS), mitochondrial calcium transport, and ROS production. This analysis also highlighted the role of uncoupling proteins (UCPs), which were downregulated in TG mice, leading to impaired mitochondrial membrane potential regulation and further exacerbation of oxidative stress. Our data confirms that UCP dysfunction contributes to ROS generation, offering a new mechanistic understanding of how LRRK2 mutations promote mitochondrial dysregulation.

Given the strong link between LRRK2 mutations, oxidative stress, and calcium dysregulation, our findings highlight several potential therapeutic strategies. First, enhancing UCP function through pharmacological activation or gene therapy may help restore mitochondrial homeostasis and reduce ROS accumulation. Second, targeting MCU to limit mitochondrial calcium overload could mitigate oxidative stress-induced damage.

Importantly, our results support the use of isradipine (an L-type calcium channel blocker) as a potential neuroprotective strategy in LRRK2-related PD (Guzman et al., 2010, 2018). While the STEADY-PD III clinical trial failed to demonstrate significant benefits in idiopathic PD, this is likely due to the intervention occurring at an irreversible stage of the disease, when over 80% of dopaminergic neurons have already been lost. The lack of efficacy does not rule out its potential in earlier stages. Future studies should focus on early intervention, particularly in high-risk individuals, such as genetically defined LRRK2-PD carriers, before significant neuronal loss occurs. Finally, Future studies should assess whether selective MCU inhibitors or UCP agonists can provide neuroprotection in LRRK2-associated PD models.

## Conclusion

Our study provides the first direct evidence that LRRK2-hR1441G mutations drive mitochondrial oxidative stress through calcium dysregulation and impaired UCP function in SNc DA neurons. We demonstrate early oxidative stress in presynaptic terminals, followed by age-dependent mitochondrial dysfunction in SNc DA neurons, highlighting a potential axon-first degeneration model in PD. Our findings suggest that targeting MCU, L-type calcium channels, and UCPs may provide novel therapeutic strategies for LRRK2-associated PD.

## Methods

### Animals

BAC LRRK2(hR1441G) transgenic (TG) mice (Stock #009604, The Jackson Laboratory) were obtained from Dr. Chenjian Li’s laboratory at Weill Medical College of Cornell University and maintained on a Taconic FVB/N genetic background. Transgenic mice expressing a roGFP construct under the tyrosine hydroxylase (TH) promoter, with a mitochondria-targeting matrix sequence, were acquired from Dr. James Surmeier’s laboratory at Northwestern University. These mice were backcrossed onto the FVB/N background for eight generations before being crossed with hR1441G mice, generating GFP-expressing hR1441G and non-transgenic (nTg) lines. All mice were maintained on an FVB/N background (Taconic).

The animals were housed in a temperature- and humidity-controlled environment with a 12-hour light/dark cycle and provided ad libitum access to food and water in a specific pathogen-free facility. All experimental procedures involving animals were conducted in accordance with the National Institutes of Health guidelines and were approved by the Institutional Animal Care and Use Committees (IACUC) of Thomas Jefferson University and The University of Georgia.

### Brain slice preparation

Mice were anesthetized using a ketamine/xylazine mixture, followed by transcardial perfusion with ice-cold, oxygenated cutting solution containing (in mM): 125 NaCl, 2.5 KCl, 26 NaHCO3, 3.7 MgSO4, 0.3 KH2PO4, and 10 glucose (pH 7.4). Following perfusion, the mice were decapitated, and the brains were immediately removed and sectioned in ice-cold, oxygenated cutting solution using a vibratome (VT1000S, Leica Microsystems, Germany). Brain slices, 250 µm in thickness, were recovered at 34°C in oxygenated artificial cerebrospinal fluid (aCSF; in mM: 125 NaCl, 2.5 KCl, 26 NaHCO3, 2.4 CaCl2, 1.3 MgSO4, 0.3 KH2PO4, 10 glucose, 2 HEPES, pH 7.4) for 30 minutes and then maintained at room temperature.

For oxygen consumption rate (OCR) measurements, 160 µm-thick slices of the striatum were prepared and transferred to respiration buffer composed of preoxygenated aCSF supplemented with 25 mM glucose, 2 mM L-glutamine, and 4 mg/mL bovine serum albumin (BSA), freshly added before use (pH 7.4, 37°C). Tissue punches were immediately taken from the slices for subsequent respiration studies.

### TH-Immunogold labeling and electron microscopy

WT and TG mice were perfused transcardially with 3.75% acrolein and 2% paraformaldehyde in 0.1 M sodium phosphate buffer (PB, pH 7.4). Brains were removed and post-fixed for 30 min in 2% paraformaldehyde before being transferred to PB. Brains were coded so investigators were blinded to genotype. Brains were sectioned into 50 µm slices in PB using a vibratome. Sections were then treated with 1% sodium borohydride in PB for 30 minutes to reduce free aldehydes, followed by extensive rinsing in PB. The tissue was subsequently transferred to 0.01 M tris-buffered saline (TBS), pH 7.6. Some sections were further processed by transferring them to a cryoprotectant solution and frozen for future use, while the remaining sections were immediately processed for immunogold-silver labeling of tyrosine hydroxylase (TH).

Blocking solution was prepared in TBS containing 3% goat serum, 1% bovine serum albumin (BSA), and 0.04% Triton X-100, and applied to the tissue for 30 minutes to enhance antibody penetration. The primary antibody, mouse anti-TH (Chemicon), was diluted 1:5000 in the blocking solution and incubated with the tissue overnight at room temperature. After incubation, sections were washed in TBS, followed by 0.01 M phosphate-buffered saline (PBS), pH 7.4. They were then incubated for 30 minutes in a washing buffer consisting of PBS, 0.8% BSA, 0.1% fish gelatin, and 3% goat serum.

Next, sections were transferred to a solution containing 0.8 nm gold-conjugated anti-mouse secondary antibody, diluted 1:50 in the washing buffer, and incubated overnight at room temperature. The following day, the sections were rinsed thoroughly in washing buffer, followed by PBS, and incubated for 10 minutes in 2.5% glutaraldehyde in PBS to stabilize the gold particles. After rinsing in PBS, sections were treated with the proprietary Enhancement Conditioning Solution (ECS; Aurion/Electron Microscopy Sciences). The sections were then incubated in Aurion R-Gent SE-EM kit reagents at room temperature for an empirically determined time of approximately 90-120 minutes. After further rinsing in ECS, the sections were transferred to PB.

For electron microscopic visualization, immunogold-silver labeled tissue was subjected to osmication, dehydration, and plastic embedding. Sections were incubated in 2% osmium tetroxide in PB for 30 minutes, followed by rinsing in PB. The tissue was then sequentially dehydrated in 30%, 50%, 70%, and 95% ethanol solutions, and subsequently exposed twice to 100% ethanol for 10 minutes each, followed by two 10-minute exposures to propylene oxide. The sections were incubated overnight in a 1:1 mixture of epoxy resin (Epon; EMBed-812, Electron Microscopy Sciences) and propylene oxide before being transferred to pure Epon for 2 hours. The sections were embedded in Epon between sheets of commercial plastic and polymerized under heavy lead weights at 62°C for 72 hours.

The embedded tissue sections containing the dorsal striatum were mounted on plastic blocks, and excess tissue was trimmed, leaving a trapezoidal region within the dorsolateral striatum. This region of interest was then sectioned into ultrathin (50-60 nm) slices using an ultramicrotome and collected onto copper-400 mesh grids. The ultrathin sections were counterstained with uranyl acetate and lead citrate.

Ultrathin tissue sections were examined using a Morgagni FEI transmission electron microscope at 36,000X magnification. Sampling of TH-immunolabeled profiles was done near the interface between tissue and epoxy resin, where maximal antibody penetration occurred. To decrease the collection of false positive results, TH-labeled axons were defined as profiles containing a minimum of three gold-silver particles, with many axons exhibiting considerably more than three particles.

### Measurement of Real-Time Oxygen Consumption Rate (OCR)

Real-time oxygen consumption rate (OCR) in acute brain slices was measured using a Seahorse XF24 analyzer (Seahorse Bioscience) with minor modifications to the manufacturer’s protocol. To optimize conditions for OCR measurement, various combinations of slice thickness (150 µm or 200 µm) and punch size (1.0 mm, 1.5 mm, and 2.0 mm) were tested. Brain slices were secured to the center of the mesh in an islet capture plate as described previously (Zhi et al., 2019).The microplate was then incubated at 37°C for 1 hour to equilibrate temperature and pH.

Drugs and inhibitors were diluted in prewarmed (37°C) respiration buffer to stock concentrations (10×, 11×, 12×, and 13× the working concentrations for ports A, B, C, and D, respectively) and preloaded into the reagent ports of a sensor cartridge. The sensor cartridge had been hydrated overnight in Seahorse calibration solution at 37°C without CO2. Each injection had a volume of 75 µL. Following a 30-minute calibration and equilibration period, OCR was measured in each well of the plate using a program consisting of 3-minute mixing, 3-minute waiting, and 2-minute measurement cycles. Baseline OCR was determined with four measurements, followed by sequential injections of port A (10 mM pyruvate, 3 X measurements), port B (20 mM Oligomycin, 8 X measurements), port C (10 mM or other concentrations of FCCP, 5X measurements), and port D (20 mM antimycin A, 6 X measurements). The parameters for load, measure, and calibration distance of probe head were set to 27,800.

### TMRM Dye labeling

Tetramethylrhodamine methyl ester (TMRM; Invitrogen) was used to monitor mitochondrial membrane potential. A 10 mM TMRM stock solution prepared in DMSO was diluted in oxygenated artificial cerebrospinal fluid (aCSF) to a final working concentration of 4 µM. Brain slices were incubated in the oxygenated TMRM-containing aCSF at 37 °C for 30 minutes to allow for dye equilibration and accumulation within the mitochondrial matrix. Following incubation, the slices were transferred to oxygenated TMRM-free aCSF and washed for 30 minutes to remove excess dye.

To assess the sensitivity of roGFP protein to oxidative stress, brain slices expressing roGFP were sequentially perfused with aCSF containing the reductant dithiothreitol (DTT, 2 mM, Sigma) for 45 minutes, followed by the oxidant aldrithiol (Ald, 100 µM, Sigma) for an additional 45 minutes. To investigate the effects of uncoupling proteins (UCPs), L-type calcium channel blockers, and antioxidants on mitochondrial membrane potential (MMP) *ex vivo*, brain slices were incubated in oxygenated aCSF containing genipin (Sigma), isradipine (Tocris-Cookson, 20 µM), or N-(2-mercaptopropionyl)-glycine (MPG, 10 µM, Sigma) for 30 minutes, respectively. To examine whether L-type calcium channel inhibition reduce or diminish calcium-dependent mitochondrial oxidant stress in the animal, we administered intraperitoneally isradipine (3 mg/kg body weight) daily for 7-10 days.

Reagents used in Seahorse experiments included bovine serum albumin (BSA) (A6003), glucose (G8270), sodium pyruvate (P2256), L-glutamine (C3126), oligomycin from streptomyces diastatochromogenes (O4876), carbonyl cyanide 4-(trifluoromethoxy) phenylhydrazone (FCCP) (C2920), antimycin A from streptomyces sp. (A8674), which were purchased from Sigma Aldrich (St. Louis, MO) and rotenone (3616, Tocris). In addition, stainless steel biopsy punches were purchased from Miltex (York, PA); and XF24 Islet Flux-Paks from Seahorse Bioscience (Billerica, MA).

### Real-Time PCR

Following the protocol described by Guzman et al., total RNA from the substantia nigra (SN) region was extracted for quantification of uncoupling protein (UCP) mRNA levels. Ventral midbrain regions, encompassing the SNc and VTA, were micro-dissected from 10-month-old LRRK2 R1441G wild-type (WT) and transgenic (TG) mice. Total RNA was extracted using TRIzol reagent (Invitrogen), and cDNA synthesis was performed via reverse transcription (Quanta Biosciences) using oligo-(dT)15 primers.

Quantitative PCR (qPCR) was carried out to analyze cDNA from each sample, employing SYBR Green as the fluorescent dye and sense/antisense primers synthesized by Integrated DNA Technologies. The qPCR cycling parameters included an initial denaturation step at 95□°C for 3 minutes, followed by 40 cycles of 15 seconds at 94□°C, 1 minute at 60□°C, and 30 seconds at 72□°C. SYBR Green fluorescence was measured after each cycle.

Relative mRNA expression levels were calculated using the comparative CT (ΔΔCT) method. Gene-specific mRNA expression was normalized to β-actin levels to obtain Δ*C*_T_ values (Δ*C*_T_ = *C*_T_(UCP4/5)□−□*C*_T_(ß-actin)). Paired Δ*C*_T_s for WT and *TG* groups and the mRNA abundance were calculated from the equation 2 ^-ΔΔ^ ^CT^. All samples were analyzed in triplicate to ensure accuracy.

### Western Blot

Mouse SNc slices were homogenized in RIPA lysis buffer containing 1X protease inhibitor cocktail and centrifuged at 21,000g for 30 minutes at 4°C. Protein concentration was determined using the BCA protein assay kit. A total of 200 µg of protein was mixed with 4X Laemmli buffer and β-mercaptoethanol to achieve a final concentration of 1 µg/µl in 1X Laemmli buffer and 10% β-mercaptoethanol, then boiled for 5 minutes at 95°C. A total of 10-20 µg of protein per lane was loaded onto 10% polyacrylamide gels and electrophoresis in Tris/glycine buffer, followed by transfer to 0.2 µm PVDF membranes. The PVDFs were blocked with 5% non-fat dry milk in TBS containing 0.05% Tween-20 (TBST) for 1-2 hours at room temperature with gentle shaking at 20 rpm, and then incubated with primary antibodies overnight at 4°C. After washing TBST three times, the PVDFs were incubated with the appropriate horseradish peroxidase (HRP)-conjugated secondary antibody for 1 hour at room temperature with gentle shaking at 20 rpm. The PVDFs were then washed three times with TBST, developed using Femto Sensitivity Substrate, and imaged using a ChemiDoc MP Imager (Bio-Rad).

### Stereotaxic injection of AAV

Mice were anesthetized with isoflurane (1-2%) or ketamine/xylazine (100 mg/10 mg/kg). A small craniotomy, approximately 0.8 mm in diameter, was made over the right midbrain (−3.4 mm caudal, +1.0 mm lateral). To achieve expression of cyto-GCaMP6 or mito-GCaMP6 in dopaminergic neurons, 0.1-0.2 µl of AAV-GCaMP6-mito virus (10^13 vg/ml, a gift from Dr. James Surmeier’s lab at Northwestern University) was pressure-injected through a syringe into the midbrain at a depth of -4.2 mm ventral from the dura surface. Following the injections, the incisional wound on the skull was sealed with tissue adhesive (3M Vetbond). The mice were placed on a heating pad until fully recovered and returned to the isolation room. Carprofen (5 mg/kg, Sigma Aldrich) was administered intraperitoneally once per day for at least 3 days to reduce pain.

### Two-photon laser scanning microscopy (2PLSM) imaging of mitochondrial functions in brain slices

The brain slices were positioned within a recording chamber and superfused with oxygenated aCSF at a flow rate of 1.5 mL/min. Optical imaging of florescent signals were acquired using an Ultima multiphoton laser scanner (Bruker) for BX51/61WI, Olympus microscope with a Titanium-sapphire laser (Chameleon-Ultra2, Coherent Laser Group), equipped with a 20 × 1.0 NA water immersion objective (XLUMPLFL20XW, Olympus). The Dendra2 protein was excited at a wavelength of 900 nm, and its fluorescence was collected at 500-560 nm. The average laser power for imaging was less than 50 mW. Images were captured in 8-bit resolution and 4 μs dwell time, with regions of interest set at 512 × 512 pixels. The sampling rate ranged from 0.2 to 0.5 seconds per frame. All experiments were conducted at 33-35 °C.

### Mitochondrial roGFP imaging

Acute brain slices with roGFP expressed in mitochondria were put and anchored under the microscope with the oxygenated aCSF perfusion at physiological temperatures (34–35 °C). The optimal excitation wavelength of roGFP signals were measured at different conditions: physiological, fully reduced and fully oxidized. And the excitation wavelength (900nm and 800nm) was chosen to excite roGFP protein with a pixel size between 0.16 and 0.23 m and a 4μs pixel dwell time. roGFP fluorescence (490–560 nm) was differentiated and collected by PMT detector. Laser intensity and PMT gain were kept consistent or adjusted in proportion. Records with a drifting baseline were excluded for data analysis.

### 2PLSM cyto-GCaMP6 and mito-GCaMP6 imaging in acute brain slices

Mitochondrial Ca2+ levels were measured with the mitochondrially targeted Ca2+-sensitive probe mito-GCaMP617. Two-three weeks after AAV virus injection, mice were prepared for coronal brain slice sectioning following transcardiac perfusion with ice-cold, oxygenated cutting solution. After a recovery period of 30-45 minutes, the coronal brain slices were placed in a chamber perfused with oxygenated aCSF. To measure free Ca2+ levels in mitochondria, the slices were sequentially treated with the following solutions: (1) regular aCSF, (2) 500 µM EGTA + aCSF without Ca2+ for 30-40 minutes, (3) 500 µM EGTA + 1 µM ionomycin + aCSF without Ca2+ for 30 minutes, and (4) 1 µM Ionomycin + aCSF with high Ca2+ (3 mM) for 20 minutes. Time series images were acquired every 5 minutes to determine the minimal and maximal fluorescence intensity. The fluorescence intensity at the end of treatment (2) was defined as the minimum fluorescence (Fmin), and the fluorescence intensity at the end of treatment (4) was defined as the maximum fluorescence (Fmax). The relative free calcium level was calculated using the equation: (F - Fmin) / (Fmax - Fmin).

### Mitochondrial TMRM imaging

Brain slices from LRRK2 hR1441G with TH-GFP expression were incubated in 4 μM TMRM for 30 min at 34–35 °C; excess dye was washed out with a TMRM-free ACSF solution. Brain slices were put in the chamber under the microscope for imaging following aCSF solution perfusion at physiological temperatures (34–35 °C). Fluorescence (550–640 nm) was collected using an 830-nm excitation beam. Time-series scanning (300 frames) was performed with a 4-μs dwell time per pixel and 2-3 frames per second scanning rate. The duration of drug applications, including MPG, genipin and isradipine, was controlled at 30 mins.

### Digital spatial profiling and scRNA sequencing

Formalin-fixed paraffin-embedded (FFPE) sections were processed and subjected to GeoMx RNA profiling (Nanostring Technologies, Seattle, WA, USA). ROIs/AOIs were defined using TH and IBA1 markers, and gene set enrichment analysis (GSEA) was performed to identify enriched pathways in LRRK2 mutant microglia. Further methodological details are detailed below.

#### FFPE Tissue Preparation

Formalin-fixed paraffin-embedded (FFPE) tissues were prepared as follows: mice were anesthetized, and brains were fixed via transcardial perfusion with 10% neutral buffered formalin (Sigma-Aldrich). Brains were subsequently removed, post-fixed in the same formalin solution for 24 hours at 4°C and immersed in 70% ethanol for at least 1 hour. Tissues were processed and embedded in paraffin using an automated tissue processor (Tissue-Tek VIP, Sakura Finetek USA Inc.) following the programmed protocol: 95% ethanol for 1h15m, 100% ethanol for 1h15m, 100% ethanol for 1h30m, 100% xylene for 1h, 100% xylene for 1h30m, 100% xylene for 1h30m, paraffin for 1h30m at 58°C, and paraffin for 2h at 58°C. FFPE tissues were then immersed in molten paraffin at 60°C in a metal mold to form FFPE blocks. These blocks were cooled at 4°C and stored at the same temperature until further use.

#### Tissue Sectioning and Slide Preparation

The midbrains of FFPE mouse brains were coronally sectioned at 5 μm using a rotary microtome. Sections containing the substantia nigra (SN) and ventral tegmental area (VTA) were mounted on Leica BOND Plus microscope slides, with each slide holding 6 sections—3 from wild-type mice and 3 from transgenic mice. The slides were submitted to the Flow Cytometry and Human Immune Monitoring Core at Thomas Jefferson University for RNA profiling via the GeoMx RNA assay (Nanostring).

#### ROI and AOI Selection

Tissue sections were stained with fluorescent morphology markers and nuclear counterstain for the identification of regions of interest (ROIs) and areas of interest (AOIs). The morphology markers included an anti-tyrosine hydroxylase (TH) antibody conjugated with Alexa 488 (dopaminergic neuron marker, Sigma #MAB5280X), an anti-GFAP antibody conjugated with Alexa 594 (astrocyte marker), and an anti-Iba1 antibody conjugated with Alexa 647 (microglia marker, Millipore #MABN92-AF647). The nucleus marker was Syto-13. For each tissue section, 2-3 ROIs were selected within the SN or VTA, and each ROI was subdivided into 1-3 AOIs representing TH+, GFAP+, or Iba1+ cell populations. Each AOI contained a minimum of 20 cells.

#### Quality Control

Raw counts from GeoMxTM Digital Spatial Profiling (DSP) were processed using GeoMxTM Software. Out of 95 segments collected from 6 mice (3 harboring hLRRK2 R1441G mutations and 3 wild-type), 11 segments were excluded due to failure in quality control (QC) analysis. Following QC, 8021 out of 20175 gene targets in the GeoMx Mouse Whole Transcriptome Atlas passed QC and were retained for downstream analyses.

#### Data Normalization and Analysis

To address systematic variability between AOIs, GeoMxTM DSP raw counts were normalized to the 75th percentile of expression for each AOI, as described in the NanoString GeoMxTM Data Analysis User Manual. Differential expression analysis was performed using a negative binomial model, implemented in the DESeq2 package in R.

#### Gene Set Enrichment Analysis (GSEA)

Gene Set Enrichment Analysis (GSEA) was carried out to identify dysregulated pathways in dopaminergic neurons between mice carrying hLRRK2 R1441G mutations and wild-type controls. GSEA was conducted using the ClusterProfiler package (version 4.12) in R, with gene sets ranging in size from 10 to 500 genes. Enrichment terms with p-values below 0.05 were considered significantly enriched. Additionally, GSEA was extended using gene set signatures from the Reactome pathway database and the KEGG dataset respectively, with normalized enrichment scores (NES) calculated to identify biological pathways linked to hLRRK2 R1441G mutations in dopaminergic neurons.

### Image Data analysis

Images were processed using open-source software Fiji (NIH) and commercial software Matlab (Version 8.5.0 R2015a, Mathworks). Registration had been taken to perform the intensity-based alignment of images at the different time point. DA terminals in the striatum and DA soma in SNc area were segmented and regarded as the regions of interest (ROIs), the ratio of fluorescence intensity in these ROIs excited by 800nm beam divided by corresponding ROIs excited by 900nm beam was used to measure the level of oxidation stress. Flickering area of TMRM labeling were selected as ROIs and the flickering frequency in the ROIs was determined by counting the number of transitions in 100-s epochs. The change in mitochondrial membrane potential during flickering was estimated from fluorescence in ROIs.

### Statistical analysis

Data are presented as mean ± standard deviation (SD) where applicable. Statistical analysis was performed with GraphPad Prism 8 (GraphPad software Inc.). Unpaired Mann-Whitney test, one-way ANOV, and two-way ANOVA, were applied, with a P-value of < 0.05, considered statistically significant.

**Table 1:** Key Source and reagent.

| Reagent or Resource | Source | Identifier |
| --- | --- | --- |
| **Reagent** |  |  |
| Tetramethylrhodamine methyl ester (TMRM) | Invitrogen | Cat #T668 |
| Dithiothreitol (DTT) | Sigma | Cat #D0632 |
| Aldrithiol | Sigma | Cat #143049 |
| Genipin | Sigma | Cat #G4796 |
| Isradipine | Tocris | Cat #2004 |
| N-(2-mercaptopropionyl)-glycine (MPG) | Sigma | Cat #M6635 |
| Bovine serum albumin (BSA) | Sigma | Cat #A6003 |
| Sodium pyruvate | Sigma | Cat #P2256 |
| L-glutamine | Sigma | Cat #C3126 |
| Oligomycin | Sigma | Cat #O4876 |
| Carbonyl cyanide 4-(trifluoromethoxy) phenylhydrazone (FCCP) | Sigma | Cat #C2920 |
| Antimycin A | Sigma | Cat #A8674 |
| Rotenone | Tocris | Cat #3616 |
| **Source** |  |  |
| XF24 Islet capture microplate | Agilent | Cat #101122-100 |
| XF24 Islet capture FluxPaks | Agilent | Cat #101174 |

**Table 2:**
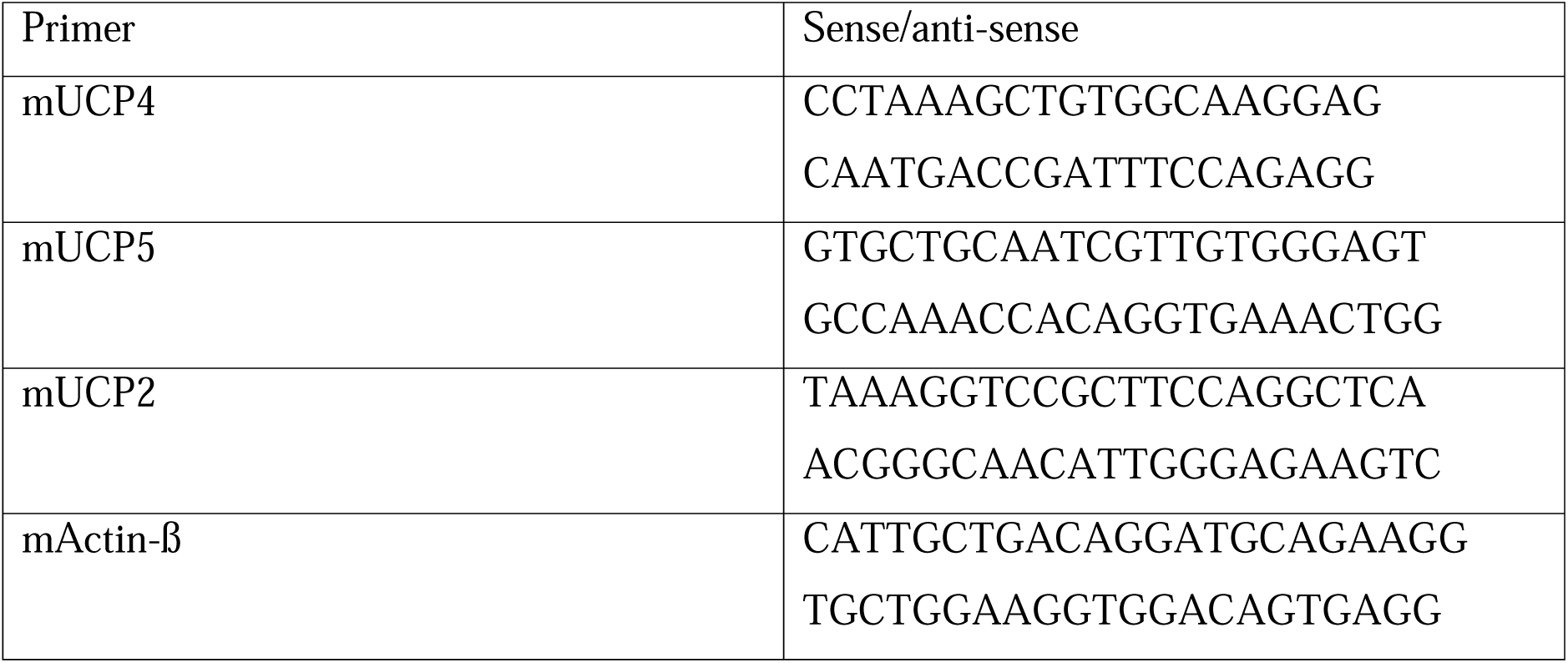
qPCR primer.

**Table 3:**
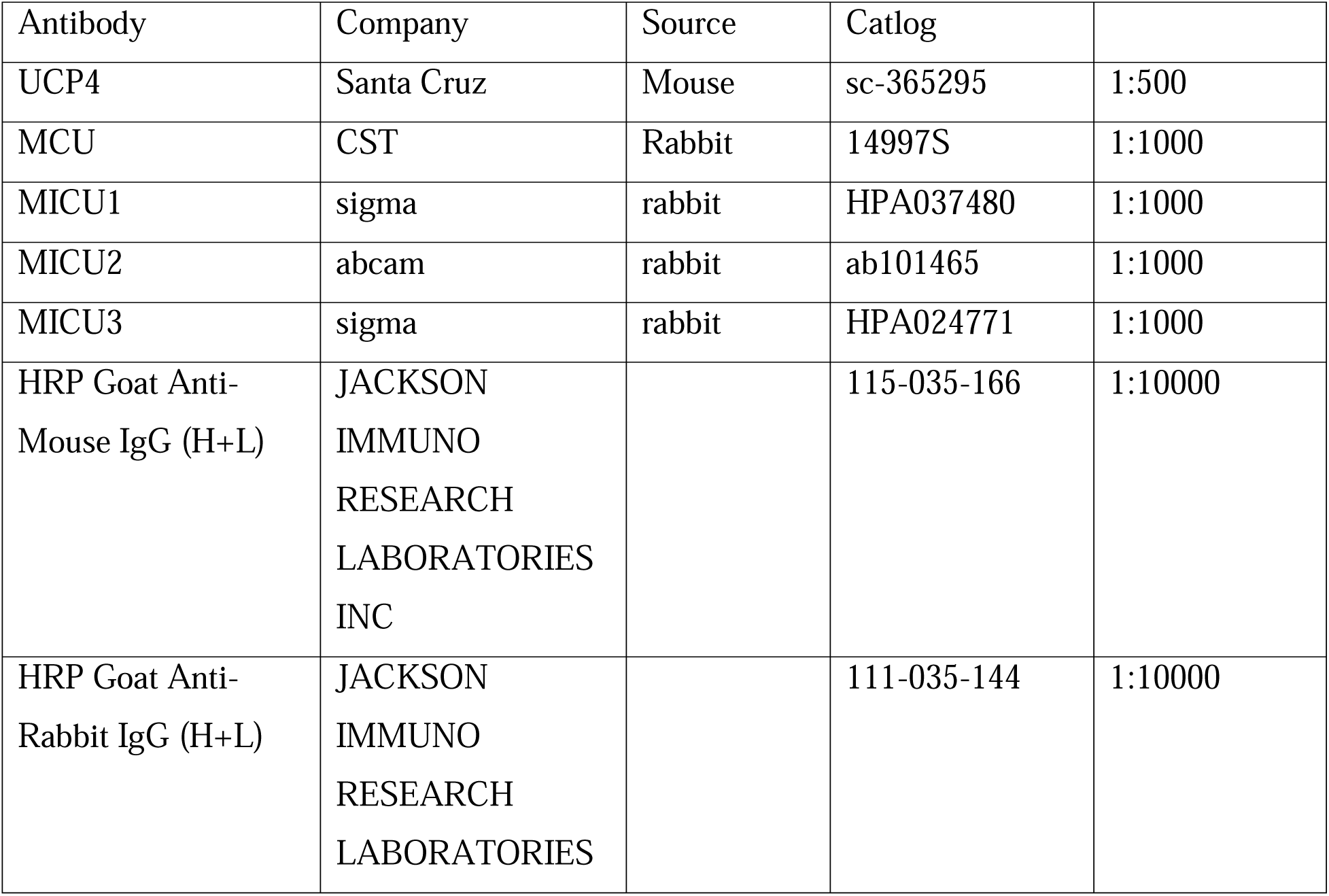
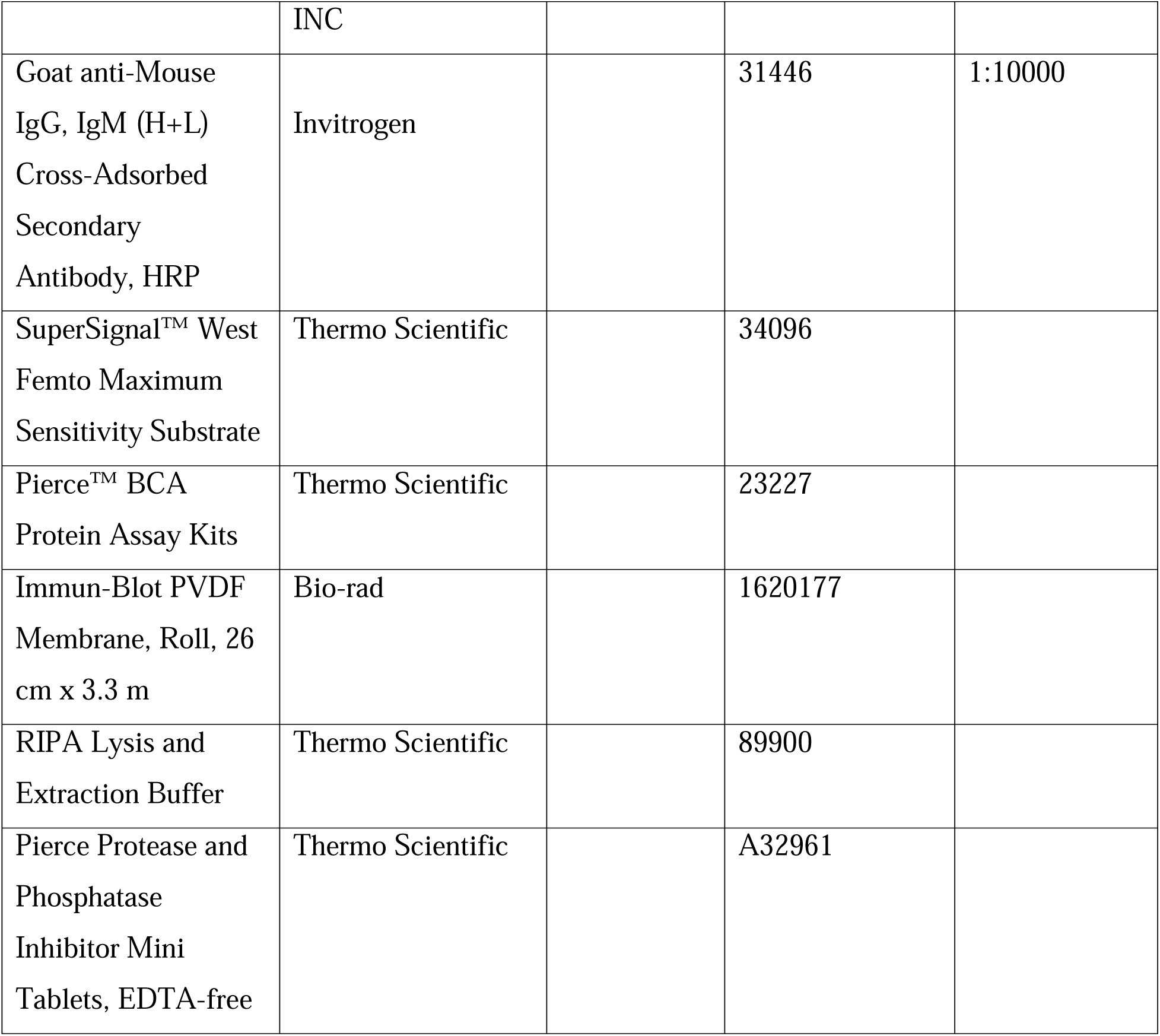
Antibody List.

## Data availability

Data is available from the corresponding authors on request.

## Acknowledgments

We thank Dr. D. James Surmeirer at Northwestern University for providing the TH-mito-roGFP mice and the AAV-TH-GCamp6 virus. We also thank Dr. Susan R. Sesack at the University of Pittsburgh who prepared tissue from wild-type and LRRK2-R1441G transgenic mice for immunogold labeling and performed photographic sampling of striatal sections under electron microscopy. This work was supported by the National Institute of Neurological Disorders and Stroke (NINDS) (Grant NS098393, NS097530 and NS128005 to H.Z.), and startup funds from Thomas Jefferson University (H.Z.) and the University of Georgia (H.Z.).

## Author contributions

H.Z. conceived of and supervised the project. Y.X.C. and H.Z. wrote the paper. Y.X.C. and H.Z. analyzed the data, with input from all authors. Y.X.C. performed 2p experiments. L.T.Z performed the Seahorse assays. L. T. Z. performed the quantitative real-time RT-PCR (qPCR). S.Q.C. performed WB.

## Supplementary material

Supplementary material is available online.

