## Supplemental Figs and Tables for "The Human LRRK2-R1441G Mutation Drives Age-Dependent Oxidative Stress and Mitochondrial Dysfunction in Dopaminergic Neurons"

**Extended Data Figures:**

**Extended Data Fig.1**: **Mitochondrial respiration capabilities in LRRK2 hR1441G mice at 3 months and 6 months of age**.

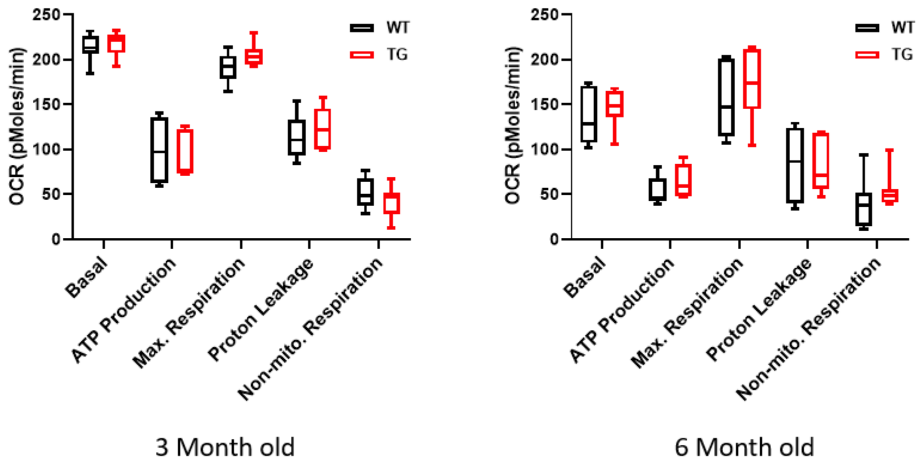


**Extended Data Fig.2**: Spectrum of roEGFP expressed in SNc dopaminergic neurons under physiological conditions (a) and after treatments with DTT and Ald, respectively (b).

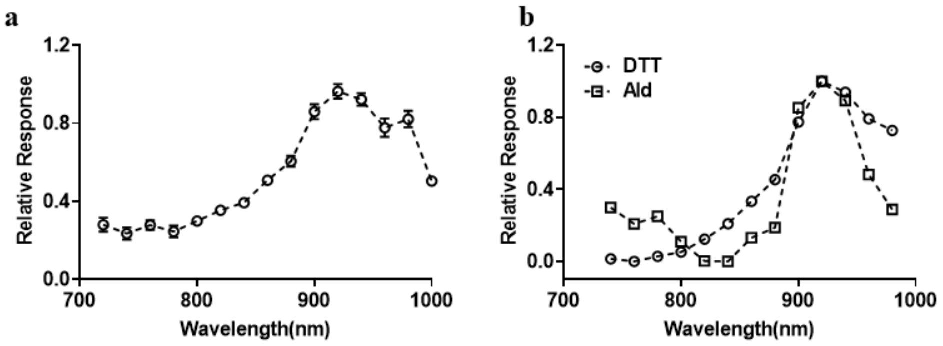

### Extended Data Fig.3: Sensitivity of roEGFP to sequential DTT and Ald treatments

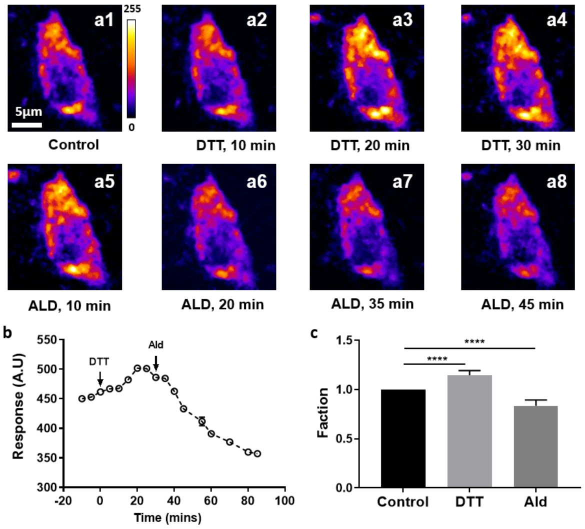


**a1–a8**, Representative images of SNc dopaminergic neurons at different time points following DTT and Ald treatments. **b**, Quantification of roEGFP fluorescence response to DTT and Ald treatments. c, Statistical analysis of roEGFP responses to sequential DTT and Ald treatments (N = 4, n = 6).

### Extended Data Fig.4: Validation of the ratiometric measuring oxidative levels

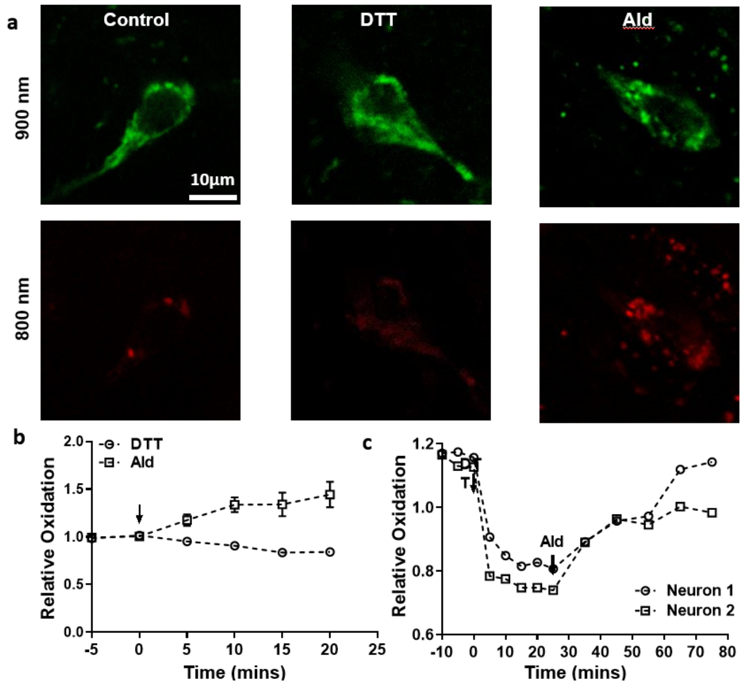


**a**, Representative roEGFP images under different conditions (left: physiological; middle: fully reduced with DTT; right: fully oxidized with Ald) captured at two excitation wavelengths (900 nm and 800 nm). **b**, Validation of the ratiometric method for measuring roEGFP response to DTT and Ald treatments individually. **c**, Validation of the ratiometric method for assessing roEGFP response to sequential DTT and Ald treatments.

**Extended Data Fig.5: Oxidative levels in SNc DA neurons in hR1441G WT and TG mice at three different ages**

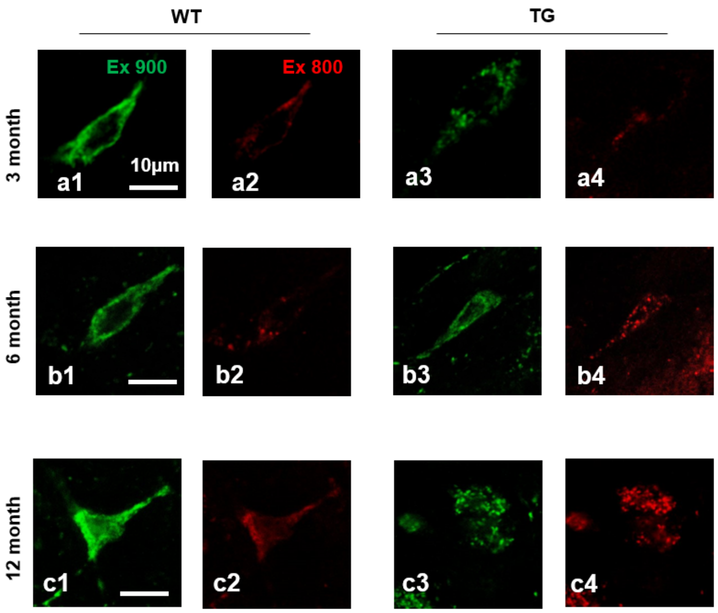


**a1-a4, b1-b4, c1-c4**, Representative images of roEGFP expressed in SNc dopaminergic (DA) neurons in LRRK2 R1441G wild-type (WT) and transgenic (TG) animal models, excited by two different wavelengths (900 nm and 800 nm), across three age groups: 3 months, 6 months, and 12 months, respectively.

**Extended Data Fig.6**: **Oxidative levels of dSTR DA terminals in hR1441G WT and TG mice at three different ages**

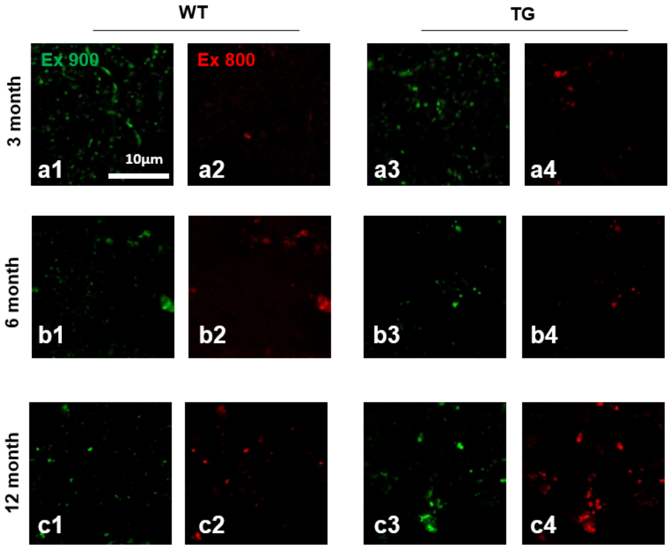


**a1-a4, b1-b4, c1-c4**, Representative images of roEGFP expression in the striatum of LRRK2 R1441G wild-type (WT) and transgenic (TG) animal models, excited by two distinct wavelengths (900 nm and 800 nm), across three age groups: 3 months, 6 months, and 12 months.

**Extended Data Fig.7**: Oxidative levels of VTA DA neuron **in hR1441G WT and TG mice at two different ages**

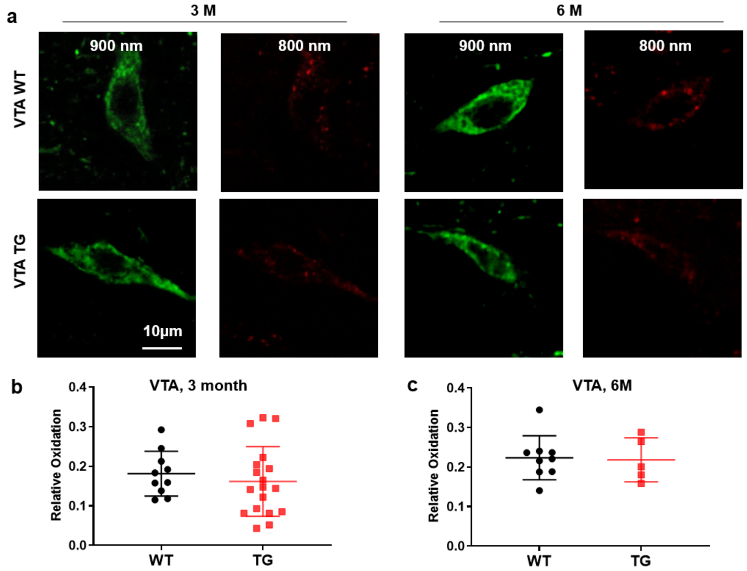


**a**, Representative roGFP images of DA neurons within VTA area at the excitation of laser beam: 900nm (Green) and 800nm (Red), between WT and TG at the different ages (3m and 6m). b, quantitation analysis of relative oxidation at 3-month-old animals. (3 pairs, DA neurons: WT, 10, TG, 18). **c**, quantitation analysis of relative oxidation at 6-month-old animal (3 pair, DA neurons: WT,10, TG, 6).

**Extended Data Fig.8**: Identification of TMRM-labeled SNc DA neurons based on TH-GFP expression

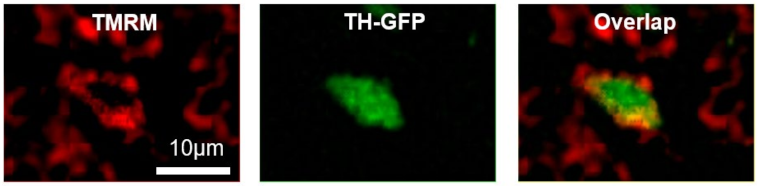

**Extended Data Fig.9**: **R1441G mutation decreased the aged-dependent flickering events in SNc DA neurons**

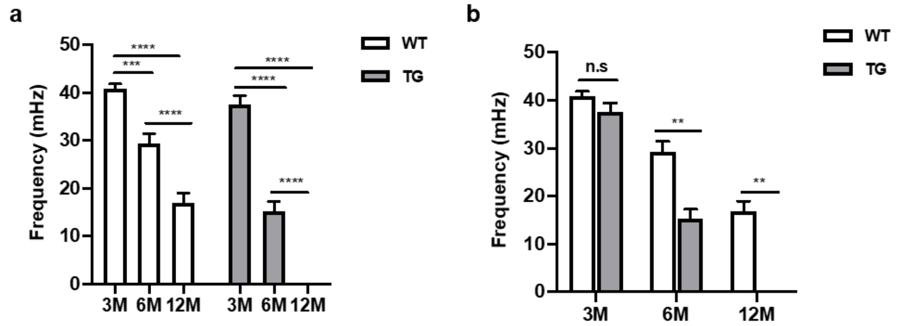


**a**, MMP flickering events are age dependent. **b**, MMP flickering frequency decreased in TG (3-month mice: 3 pair, DA neuron, WT 6, TG, 6; 6-month mice: 4 pair, DA neurons: WT 8, TG, 8; 12-month mice:  3 pair, DA neurons: WT, 7, TG, 6).

**Extended Data Fig.10**: Enrichment analysis of differentially expressed gene (DEG) between hLRRK2 R1441G and Wildtype in KEGG gene set

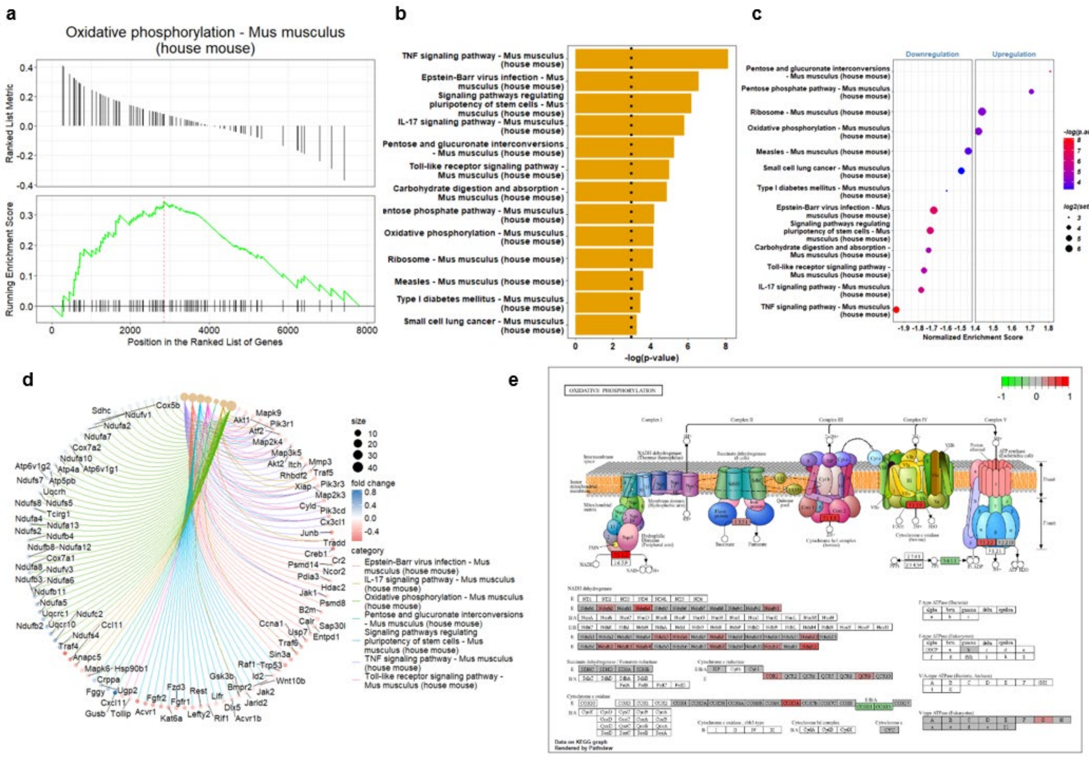


**a,** Enrichment plot comparing hLRRK2 R1441G to Wildtype in the KEGG gene set. The peak in the plot reflects the upregulation of gene sets associated with the enrichment term: Oxidative phosphorylation – Mus musculus (house mouse). **b**, Reactome pathway analysis of differentially expressed genes (DEGs) between hLRRK2 hR1441G and Wildtype. **c**, Gene set enrichment analysis (GSEA) of the Reactome gene set based on DEGs from hLRRK2 R1441G versus Wildtype, represented as a hierarchically clustered heatmap. The color scale corresponds to the -Log$_{10}$ of the p-value for the enrichment score, and the size of the circles corresponds to the number of leading genes in each enriched term. **d**, CNET plot from gene set enrichment analysis, illustrating the complex associations between genes and significantly enriched terms. Node size is proportional to the significance of the enrichment (adjusted p-value), while the color of the gene nodes represents fold change values. Edges indicate the associations between genes and the enriched terms. **e**, Visualization of the Oxidative phosphorylation enrichment term, highlighting upregulated and downregulated genes.

### Extended Data Video Legends

**Extended Data Video 1**: Representative movies of mitochondrial membrane potential of SNc DA neuron between WT (A) and TG (B).

**Extended Data Video 2**: MPG treatment reduced the MMP flickering frequency of SNc DA neurons.

**Extended Data Video 3**: L-type calcium channel blocker, Isradipine, reduced the MMP flickering frequency of SNc DA neurons.

**Extended Data Video 4**: Genipin reduced the MMP flickering frequency of SNc DA neurons.
